# Cotton Microbiome Profiling and Cotton Leaf Curl Disease (CLCuD) Suppression through Microbial Consortia associated with *Gossypium arboreum*

**DOI:** 10.1101/2023.07.12.548745

**Authors:** Rhea Aqueel, Ayesha Badar, Nazish Roy, Qandeel Mushtaq, Aimen Fatima Ali, Aftab Bashir, Umer Zeeshan Ijaz, Kauser Abdulla Malik

## Abstract

The failure of breeding strategies has caused scientists to shift to other means where the new approach involves exploring the microbiome to modulate plant defense mechanisms against CLCuD. The cotton microbiome of CLCuD-resistant varieties may harbor a multitude of bacterial genera that significantly contribute to disease resistance and provide information on metabolic pathways that differ between the susceptible and resistant varieties. The current study aimed to explore the microbiome of CLCuD-susceptible *Gossypium hirsutum* and CLCuD-resistant *Gossypium arboreum*. Microbial community surveys performed using 16S rRNA gene amplification revealed that *Pseudomonas* inhabited the rhizosphere while *Bacillus* was predominantly found in the phyllosphere of CLCuV-tolerant *G. arboreum*. The study was done for the leaf endophyte, leaf epiphyte, rhizosphere, and root endophyte of the two cotton species. Furthermore, our disease incidence assay using pot experiments has revealed mechanistic insights through salicylic acid-producing *Serratia spp*. and *Fictibacillus spp*. isolated from CLCuD-resistant *G. arboreum*, which exhibited viral disease suppression and induced systemic resistance in CLCuD susceptible *G. hirsutum*.

## Introduction

Cotton Leaf Curl Disease (CLCuD), transmitted by the whitefly *Bemisia tabaci*, has devastated Pakistan’s cotton crop for the past three decades [1, 2]. Insect pests account for 37% of cotton yield losses, whereas *Bemisia tabaci* is responsible for 50% of the total loss in cotton production due to insects [3, 4]. A few CLCuD-tolerant lines of *Gossypium arboreum* have been developed through conventional breeding approaches, but the worldwide extensively cultivated *Gossypium hirsutum* remains highly susceptible to CLCuD [5, 6].

Plant-associated microbial communities reveal much about the correlation between microbiome composition and functional profiles [7]. The potential of the rhizosphere microbiome in suppressing fungal and bacterial diseases has been tapped [8, 9], but the plant microbiome’s role in suppressing viral diseases is still unexplored [10]. The plant microbiome functions as one of the key determinants of plant health and productivity by providing defense against environmental stress and pathogens. Biotic or abiotic factors affecting plant-microbe interactions contribute to distinct microbial communities spread in the three plant regions, namely rhizosphere, phyllosphere, and endosphere [11, 12]. The microbiome of cotton plants is highly affected by their genotype and developmental stages. Each cultivar’s production of genotype-specific root exudates recruits specific types of microbial communities. Similarly, at various developmental stages, different plant hormones recruit specific microbes [13]. Recent studies suggest that variation in plant genotypes, even at a single gene locus, can significantly impact rhizosphere microbiomes [14, 15].

Plant phyllosphere microbiotas play a crucial role in activating plants’ antioxidant status by producing pathogenesis-related (PR) proteins and demonstrate their capability as a promising approach for disease control management and induction of systemic resistance [16-19]. At the phylum level, the phyllosphere bacterial communities typically include Actinobacteria, Bacteroidetes, Firmicutes, and Proteobacteria, dominated by Alpha proteobacteria and Gamma proteobacteria [20, 21]. Plant secondary metabolites such as glucosinolates influence bacterial colonization in the phyllosphere and exhibit plant pathogen-inhibiting potential [22]. The phytohormone Salicylic Acid (SA) has also been studied as a protective regime against CLCuD in cotton plants where less incidence of disease and increased PR protein expression were reported [23]. The SA signaling pathway is critical to immunity against biotrophic pathogens [24]. The endosphere-associated microbes aid in plant growth and development by modulating pathways involved in plant defense and metabolism. These metabolic interactions are crucial in enhancing nutrient uptake and providing plant tolerance against biotic and abiotic stresses [25, 26].

Using the sensitive approach of next-generation sequencing-based metagenomics, we conducted complex microbial community analyses of the two cotton species, *Gossypium hirsutum*, and *Gossypium arboreum*, under CLCuD attack. We aim to highlight the role of the microbiome in suppressing plant viral disease, which has yet to be elucidated. The microbiome’s diversity reflects the correlation between microbial composition and metabolic pathways playing key roles in the cotton plant under Cotton Leaf Curl Virus (CLCuV) infection.

## Materials and Methods

### 2.1. Study Site and Sample Collection

Sampling was carried out at Four Brothers Research Farm and the Greenhouse at Forman Christian College University, Lahore (31.5204° N, 74.3587° E). The region has a semi-arid climate with annual temperatures ranging from 13.5-40°C and an average of 628.8 mm yearly rainfall. A total of 15 cotton plants were sampled across two CLCuD susceptible and tolerant cotton species, among which there were three varieties: *Gossypium hirsutum* varieties were PFV-2 (susceptible), PFV-1 (partially tolerant), and the selected *Gossypium arboreum* variety was FDH-228 (tolerant). Plants were sampled at the vegetative stage (50 days after germination DAG). A total of five replicates were taken for each of the four selected plant compartments [Leaf Epiphyte (LE), Leaf eNdophyte (LN), RhiZosphere (RZ), Root eNdophyte (RN)]. Plants were grown under viruliferous whiteflies (containing CLCuV) infestation and exhibited disease symptoms in susceptible variety after 25 days post-infestation. CLCuD-infected leaf tissues were taken from both CLCuD infected and control plants and were subjected to DNA extraction and beta-satellite amplification for CLCuV confirmation (Supplementary Figure S1).

### 2.2. Sample Preparation & Metagenomic DNA Extraction

Rhizospheric soil (up to 3mm around root area) and roots from CLCuD are susceptible, partially tolerant, and tolerant cotton plants were collected by gently shaking the roots of the plant to get rid of the bulk soil, and roots were stored in a 50 mL falcon tube. For leaf sample collection, leaves from the upper canopy showing CLCuD symptoms were taken for susceptible varieties. For control plants, upper canopy leaves were sampled under a heavy whitefly attack to ensure CLCuV infestation. Samples were placed in an ice box until they were brought to the lab, and the standard protocol was followed for preparation before DNA extraction. For leaf epiphytes, the leaves were washed three times with 1X TE buffer containing 0.2% Triton X. The wash was collected and filtered through 0.2 µM sterile filter paper, which was used for DNA extraction. The leaves were washed with 70% ethanol, then with 3% bleach, and thoroughly with sterile distilled water (SDW) for leaf endophytes. The leaf was ground using a pestle and mortar in PBS buffer and collected in a falcon tube. Roots were sonicated in PBS buffer for 5 mins for rhizospheric soil to separate the closely adhered soil. The root was removed and used for DNA extraction of root endophytes. First, roots were washed with 70% ethanol, then 3% bleach, and several washings were given with SDW. The sterilized root was macerated in PBS buffer using a pestle and mortar and was collected in a falcon tube. DNA for all four compartments (LE, LN, RZ, RN) was extracted using the FastDNA Spin Kit for Soil (MP Biomedicals) following the manufacturer’s guidelines. Samples were homogenized in the FastPrep instrument for 40 s at a speed setting 6.0. The DNA was eluted in 30 µL of elution buffer.

### 2.3. 16S rRNA V3-V4 Amplification & Sequencing

Metagenomic DNA samples were sequenced on an Illumina MiSeq platform (Macrogen, Inc. Seoul, South Korea) using the primer set listed in Supplementary Table 1 with added Illumina adapter overhang nucleotide sequences for amplification of 16S rRNA hypervariable region V3-V4. Each reaction was performed in 25 μL total volume with 12.5 μL KAPA HiFi HotStart ReadyMix (Roche), one μL from 10 μM of each primer, one μL of each mPNA and pPNA blockers, two μL of metagenomic DNA template (10 ng/μL), and remaining volume was made up with nuclease-free water. The PCR program was set as follows 95 °C for 5 min (initial denaturation), followed by 35 cycles of 94 °C for 1 min (denaturation), 55 °C for 1 min (annealing), 72 °C for 1 min, 30 sec (extension) with a final extension of 10 min at 72 °C. PCR reactions were cleaned up with AMPure® XP beads [27].

### 2.4 Bioinformatics & Statistical Analysis

The 16S rRNA gene sequences for n = 59 samples were processed with the open-source bioinformatics pipeline QIIME2. The Deblur algorithm within the QIIME2 platform was used to recover 38,120 Amplicon Sequence Variants (ASVs). PICRUSt2 algorithm as a QIIME2 plugin was used on the ASVs to predict the functional abundance of microbial communities associated with the susceptible and tolerant cotton microbiome. Statistical analysis was performed using R software. In-depth details of bioinformatics and statistical analysis are provided in Supplementary_material.pdf.

### 2.5. Isolation and characterization of culturable microorganisms

Microorganisms from all four plant compartments (LE, LN, RZ, RN) from the three selected cotton varieties were isolated on TSA and R2A media, followed by the plate count technique. Gram Staining and biochemical characterization of bacterial isolates were done using the Quick test strips (QTS) 24 bacterial identification kits (DESTO Laboratories, Karachi, Pakistan), details of which are provided in the supplementary_file.pdf.

### 2.6. Screening bacterial isolates for phytohormone production

IAA production was estimated by colorimetric method using the Salkowski reagent [28]. The bacterial strains were grown in Luria Bertani (LB) broth containing tryptophan (100mg/10ml) for 7-10 days. Then, the cells were harvested at 10,000 rpm for 15 minutes. The pellet was discarded. Salkowski’s reagent (4 ml) was added to 2 ml of sample and was incubated at 28°C for 30 minutes. The absorbance for samples was noted in triplicates at 530nm. The sample preparation was carried out for each bacterial isolate for salicylic acid production assessment using HPLC analysis [29]. Each bacterial strain was seeded into a plastic tube containing King’s B Broth (20 ml) from a single colony on a TSA plate grown for 24h at 28°C. SA was extracted after 48h from the stationary phase. At that time, bacterial cultures contain 1010-1011 cells. The liquid culture was centrifuged at 2800g for 20 mins at 4°C. The supernatant was acidified to pH=2 with 1N HCl. The solution was filtered through 0.22 µm nylon membrane, partitioned twice with 2 ml CHCl3, and dried under a nitrogen stream at 4°C. These samples were subjected to HPLC, carried out on a WatersTM HPLC System where molecular-grade salicylic acid was used as a standard. Methanol in sodium acetate buffer was used as a mobile phase.

### 2.7. 16S rRNA Identification of SA-producing bacterial strains

Bacterial colonies were directly used for amplification of the V3-V4 region using the primer pair listed in Supplementary Table 1. The bacterial colony was directly placed in the PCR mixture. The PCR profile used for amplification of V3-V4 region: 95°C for 5min, 35 cycles of 94°C for 1min, 55°C for 1min, 72°C for 1min30sec and final extension of 72°C for 10min. Water was used as a negative control. PCR products were analyzed on 2% agarose gel. Amplified PCR products of the salicylic-producing strains were sequenced using the commercial service of Sanger Sequencing at Macrogen (Seoul, Korea). The strains were identified using the V3-V4 sequence of the 16S rRNA gene using BLAST search on DDBJ / NCBI servers.

### 2.8. CLCuD Disease Incidence Assay

#### Determination of Soil Properties

Before the pot experiments, the soil used in the study was sent to PCSIR Laboratories, Pakistan, for soil texture and chemical analysis. The following parameters were determined: pH, EC (µScm-1), organic matter, N, P, K, C, Mg, Na, and Mn, with results in Supplementary Table S5. Autoclaved soil was used for pot experiments in the climate control room.

#### Pot Experiment

The disease incidence assay was carried out at Forman Christian College (A Chartered University), Lahore, Pakistan. The three SA-producing candidate strains (R1, R2, R3) were selected for application to the CLCuD susceptible (PFV-2) and partially tolerant (PFV-1) *Gossypium hirsutum* varieties in monoassociation as well as a synthetic consortium. For each genotype, a seven-day-old seedling was inoculated with bacterial suspension/synthetic consortium at 1 × 10^8^ cfu/g via soil drench application.

Exogenous SA was applied as a foliar spray for the positive control group. The application was carried out in the climate control room. Plants were shifted to the greenhouse at 21 days post application (28 DAG) for viral inoculation. For each treatment group, a total of 30 replicates were taken. Data was collected for percentage disease and disease severity index calculation till 60 days post inoculation (DPI) [30]. The formula listed below was used for calculation. In the disease severity index, 6 is the highest disease score.

Percentage disease (%) = Diseased leaves/Total leaves x 100

Disease Severity Index (DSI) = Average Percentage Disease/ 100 × 6

#### Disease Index Statistics

Disease index (DI) statistics were based on a scale of 0–6.0 (Supplementary Table S2). In the disease severity index, 6 is the highest disease score. The average percentage disease and disease severity index were calculated using the following formulae:

Percentage disease (%) = Diseased leaves/Total leaves x 100

Disease Severity Index (DSI) = Average Percentage Disease/ 100 × 6

#### Statistical analysis

SPSS software was used for data analysis, and where Tukey’s HSD post hoc test was performed with a significance level (p < 0.05).

## Results

### Comparative microbiome analysis shows significant variability in microbial community structure based on cotton species, varieties, and compartments

Through Illumina MiSeq sequencing, 38,120 ASVs were identified among 59 samples for the phyllospheric and rhizospheric bacterial community of CLCuV-infected susceptible and resistant cotton plants. The composition of bacterial phyla and the predominant genera of the different compartments of CLCuD susceptible and tolerant cotton species can be seen in Figure 1. Analysis of four compartments (LE, LN, RZ, RN) of *Gossypium hirsutum* and *Gossypium arboreum* at phyla level revealed the abundance of top 5 abundant phyla as Proteobacteria, Firmicutes, Actinobacteria, Bacteroidota, Planctomycetota (Figure 1-C). The phyla dominating *the G. hirsutum* phyllosphere were Proteobacteria, Firmicutes, Actinobacteria, and Bacteroidota. *G. arboreum* phyllosphere had Proteobacteria and Firmicutes as dominant phyla, but it also had other phyla like Gemmatimonadota and Planctomycetota, which were absent in the *G. hirsutum* phyllosphere. Planctomycetota was among the dominant phyla in the RZ of both *G. hirsutum* and *G. arboreum*, others being Myxococcota, Verrucomicrobiota, and Deinococcota. The RN of *G. hirsutum* was dominated by Proteobacteria and Actinobacteriota, whereas in *G. arboreum*, Actinobacteriota was not that dominant. The most abundant bacterial genera in CLCuD susceptible and tolerant cotton species are depicted in Figure 1-C. *Aureimonas* and *Massilia* were the predominant genera in the phyllosphere of *G. hirsutum*. However, *Candidatus portiere* was only observed in the LN region. The RZ of *G. hirsutum* included *Caulobacteraceae, Bacillus, and Escherichia-Shigella*, whereas, *Steroidobacter* and *Streptomyces* were found in the RN only. *Caulobacteraceae* and *Acinetobacter* were present in all four compartments of *G. arboreum. Planomicrobium* was in the LE region only. *Pseudomonas* was dominant in the RZ. *Ammoniphilus, Anoxybacillus*, and *Luteolibacter* were abundant in the RN region of *G. arboreum only*.

**Figure 1.**
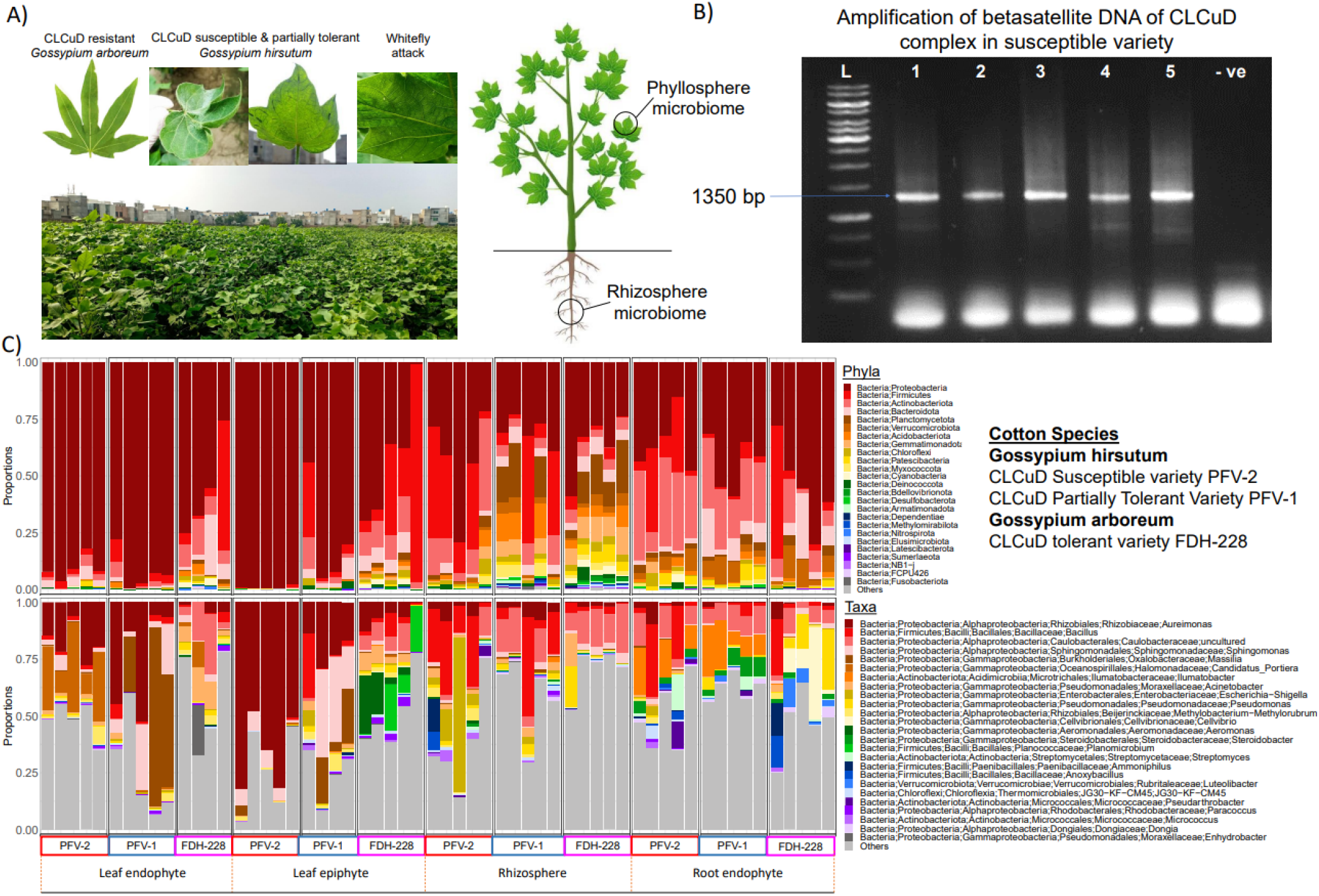
Microbiome Profiles of CLCuD Susceptible, Partially Tolerant and Tolerant Cotton Varieties. (A)-CLCuD infected cotton plants (B) Gel Image showing beta-satellite DNA amplification for CLCuV confirmation in cotton plants (C) Taxonomy Summary at Phylum and Genus Level for *Gossypium hirsutum* and *Gossypium arboreum* infected with CLCuV

Shannon and Richness, alpha diversity measures, were used for taxonomic composition. Richness is the number of ASVs/species when samples are rarefied (Figure 2-A). The highest richness in taxonomy was observed in the RZ of the CLCuD-tolerant FDH-228. The phyllosphere of CLCuD susceptible PFV-2 exhibited the lowest richness. However, the partially tolerant variety PFV-1 within *Gossypium hirsutum* species was higher than PFV-2 itself. The observed ASVs, Shannon, and richness of the bacterial communities showed significant differences between phyllospheric and rhizospheric bacteria of the three cotton varieties. Richness in metabolic functions (Figure 2-B) was higher in *Gossypium hirsutum* than *in Gossypium arboreum*. Significant results were obtained for *G. hirsutum* LN and *G. hirsutum* RN in comparison with *G. arboreum* RZ, respectively. Based on taxonomic abundance, the most balanced microbial communities belonged to the RZ of both susceptible and tolerant cotton species (Figure 2-A). In terms of functions, all compartments depicted balance, with *G. arboreum* LN and *G. hirsutum* RN being the highest (Figure 2-B). Sample dissimilarity was depicted using beta diversity measures, including Bray-Curtis distance for compositional changes (Figure 2-C), Unweighted Unifrac distance for phylogenetic changes (Figure 2-D), and Hierarchical Meta-Storms for changes in metabolic functions (Figure 2-E). It can be observed that among *Gossypium hirsutum* LE, there are two distinct groups clustered far apart of the susceptible (PFV-2) and partially tolerant (PFV-1) varieties in terms of taxa and phylogeny. Permutational multivariate analysis of variance (PERMANOVA) analysis revealed significant variability in compartments, with 28% variability in function, 14% in composition, and 8% in phylogeny. Among cotton species, the highest source of variability was in terms of function (12%) as compared to composition (4%) and phylogeny (3%). For the three cotton varieties, 3% variability was observed in composition and functions, respectively, while 2% variability was observed in phylogeny. Unweighted Unifrac and Hierarchical Meta-storms, species, and compartments significantly contributed to the beta diversity among samples.

**Figure 2.**
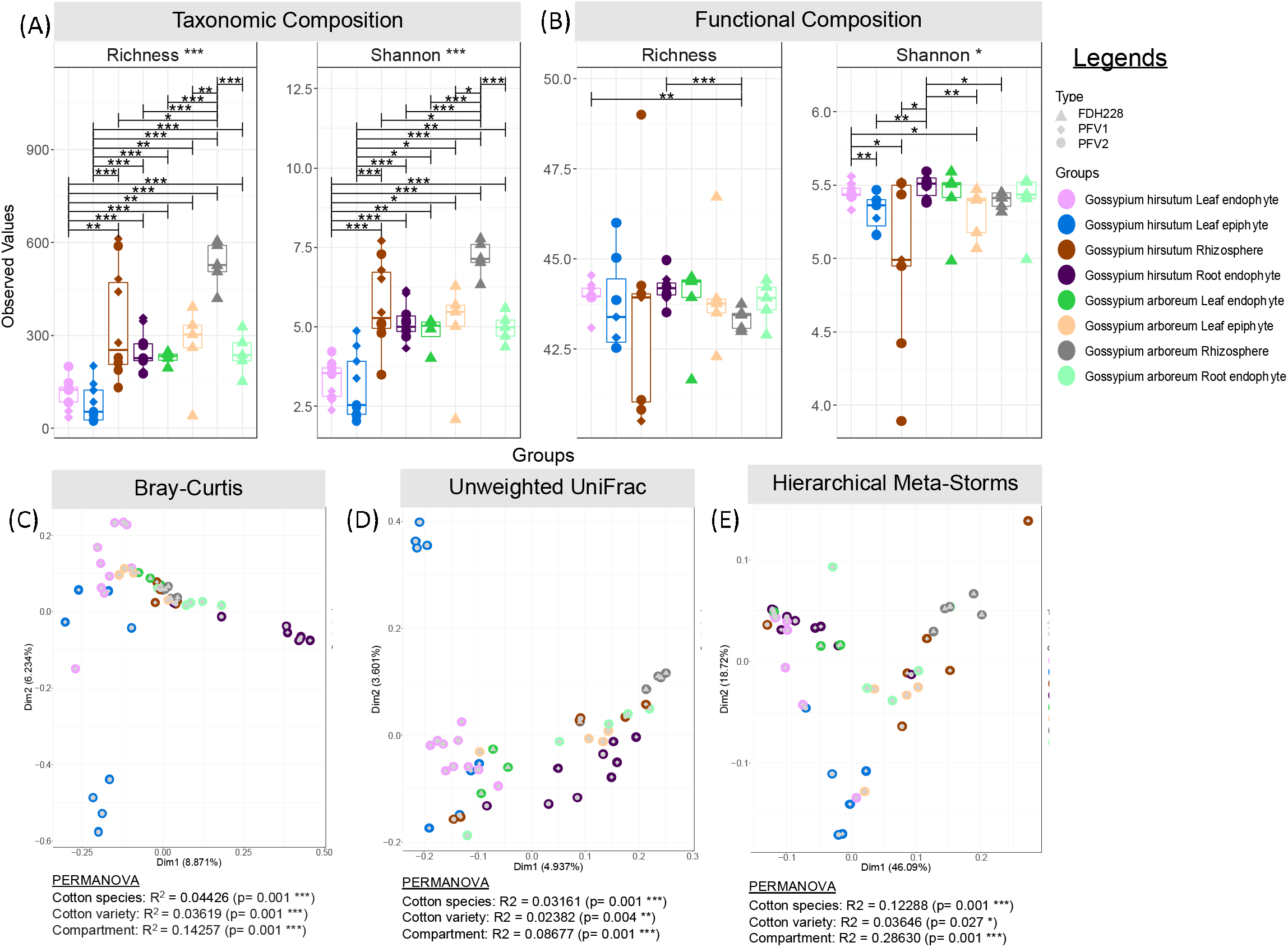
Alpha and Beta diversity results for the cotton microbiome under CLCuD attack. 1) Alpha diversity indices for A) Microbial composition; and B) MetaCyc pathways returned from PICRUSt2 procedure. Lines for panels A and B connect two sample groups at statistically significant levels indicated by asterisks as * (p < 0.05), **(p < 0.01) or ***(p < 0.001). 2) Beta diversity represented through principal coordinate analysis (PCoA) plots calculated using Bray-Curtis (A), unweighted UniFrac (B), and hierarchical meta-storms (C) distances. The ellipses represent the 95% confidence interval of the standard error of the ordination points of a given grouping. Beneath the figures, PERMANOVA R^2^ explains significant variability in microbial community structure based on Cotton species, varieties, and compartments.

### The core microbiome of the CLCuD susceptible and tolerant cotton species identified at the genus level

A core microbiome represents a set of microbes that persists in a host’s internal or surrounding environment despite location or developmental stage changes. In the CLCuD susceptible variety PFV-2 (Supplementary Figure S2), the bacterial core of LE contained 17 ASVs with *Aureimonas, Sphingomonas, Methylorubrum, Bacillus*, and *Massilia* as the most prevalent genera. *Candidatus portiere, Aureimonas, Arsenophonus, Sphingomonas*, and *Methylorubrum* dominated the LN region out of 31 ASVs. The RZ contained 26 ASVs, with *Bacillus, Pseudarthrobacter, Azospirillales, Shigella*, and *Acinetobacter* highlighted as the most prevalent. Of 58 ASVs, *Alphaproteobacteria, Bacillus, Ilumatobacter, Steroidobacter*, and *Streptomyces* were predominantly present in the RN of PFV-2. The LE, LN, and RZ of CLCuD tolerant FDH-228 have a high abundance core microbiome compared to PFV-1 and PFV-2. Overall, the bacterial core of FDH-228 (Supplementary Figure S4) contained 96 ASVs (RN) with *Vicinamibacterales, Pseudomonas, Sphingomonas, Cellvibrio*, and *Bacillus* as the most predominant genera. The LN compartment (44 ASVs) had *Acinetobacter, Caulobacteraceae, Aureimonas, Methylobacterium-Methylorubrum*, and *Pseudomonas* in prevalence. Of an abundant 224 ASVs in the RZ, the genera *Micavibrionaceae, S0134_terrestrial group, Sphingomonas, MND1*, and *Bacillus* were the most dominant. In the tolerant species, *Aureimonas, Caulobacteraceae, Aeromonas, Bacillus*, and *Acinetobacter* were the top abundant bacterial genera among 145 ASVs (LE). Only the RN compartment had a higher abundance core microbiome in partially tolerant PFV-1 (166 ASVs) as compared to PFV-2 (58 ASVs) and FDH-228 (96 ASVs).

### Distinct subsets of bacterial species in CLCuV-infected cotton plant rhizosphere and phyllosphere compartments

The association between the selected cotton plant varieties and compartments with the minimal subset of microbes was studied using the CODA LASSO variable selection approach. Positive or negative associated microbes were identified with the covariate of interest. The bacterial ASV discriminant in the RZ of PFV-1 and FDH-228 are shown in Supplementary Figure S5 (A), where *Methylophaga* and *Candidatus adlerbacteria* increased in FDH-228. In the LE compartment (Supplementary Figure S6), *Aeromonas* ASVs were positively associated with the CLCuD-resistant FDH-228, while they were negatively associated with the partially resistant PFV-1. On the contrary, *Thermus* was positively associated with the partially resistant variety PFV-1 and showed a negative association with the resistant FDH-228 in the LE region. In the RZ compartment (Supplementary Figure S8), *Ahniella* and *Nocardia* were positively associated with the CLCuD susceptible variety PFV-2 while showing a negative association with the resistant FDH-228. *Gemmatimonas* were positively associated with FDH-228 in the RZ and negatively associated with the susceptible variety. The significant positive and negative associations of bacterial ASVs associated with different plant compartments among the three cotton varieties have been shown in Supplementary Figures S5 – S11. The top 25 genera having significant correlations have been listed in Supplementary Table S3.

### Metabolic pathways abundance in CLCuV-infected cotton plant rhizosphere and phyllosphere

To elucidate the functional disparity between CLCuD susceptible and tolerant cotton species, a total of 197 MetaCyc pathways were predicted, including cellular processes, amino acid metabolism, carbohydrate metabolism, and resistance mechanisms, using the PICRUSt2 algorithm. The MetaCyc pathways of the archaea, such as those involved in phosphopantothenate, flavin, and chorismate biosynthesis, are positively associated with the *Gossypium arboreum* RN microbiome as compared to the partially tolerant PFV-1 (Supplementary Figure S12). Pathways such as formaldehyde assimilation by methanogenic bacteria and phosphopanthothenate biosynthesis by archaea were positively associated with FDH-228 and negatively associated with PFV-2 in the RZ microbiome (Supplementary Figure S13). The super pathway of salicylate degradation depicts a positive association with the RZ microbiome of susceptible PFV-2, whereas a negative association with tolerant FDH-228. Ectoine, Palmitate, and Sucrose biosynthesis pathways were positively associated with the LN microbiome of susceptible PFV-2 as compared to tolerant FDH-228 (Supplementary Figure S14). All detailed pathways showing differential abundance are shown in Supplementary Figures S12-S17.

### Key genera associated with sources of variability

To find the positive or negative association between microbial communities and the sources of variation, we have fitted a regression model, *the Generalised Linear Latent Variable Model* (GLLVM), which also gives us correlations between microbial species after accounting for the confounders (Cotton Species, Cotton Variety, CLCuV Susceptibility, and Compartment) as given in Supplementary Figure S18. Most of the microbes were positively associated with the RZ and RN plant compartments with reference to LN. The below-ground interactions and possibly root exudate chemistry are major contributing factors to these positive associations with the environmental covariates understudy. Focusing on the SA-producing bacteria, we found all of them: *Sphingomonas, Bacillus, Pseudomonas, Nocardioides*, and *Paenibacillus* to be positively associated with CLCuV resistance. Only *Sphingomonas* was positively associated with the LE in *Gossypium hirsutum. Bacillus, Pseudomonas*, and *Nocardioides* are positively associated with the RZ and the partially tolerant variety PFV-1.

### The microbiome of *Gossypium arboreum* mediates viral disease suppression in *Gossypium hirsutum*

From all three varieties and all four compartments, a total of 30 bacterial isolates were characterized for their SA production (details given in Supplementary Table S4), three of the SA-producing genera (some mentioned above) were identified from the phyllosphere of CLCuD-resistant FDH228 with the potential to see their utility as biocontrol agents. These are shown in Figure 5A and are uncultured *Serratia spp*., *Bacillus spp*., and *Fictibacillus spp*. Through pot experiment, our results exhibit the efficacy of these SA-producing bacterial strains, particularly uncultured *Serratia spp*., as a biocontrol agent on the partially tolerant variety PFV-1 of *Gossypium hirsutum*. In the PFV1 variety, the plants applied with *Serratia spp*. No CLCuD symptoms were developed at 20 days post inoculation (DPI). The control group of PFV1, on the other hand, displayed severe leaf curling and the dwarfing of the plant at 20 DPI (Figure 5-B & 5-C). The average percentage of disease in the susceptible variety PFV-2 is shown in Figure 5D, where the SA-producing strain, R1 uncultured *Serratia spp*., exhibited less than 10% CLCuD progression as compared to the PFV2.nA control group (25% at ∼35 to 40 DPI). The percentage of disease in the partially tolerant variety PFV-1 is shown in Figure 5E. The group treated with SA-producing strain R1 uncultured *Serratia spp*. and SA-producing strain R3 *Fictibacillus spp*. exhibited less than 10% CLCuD progression as compared to the PFV1.nA control group (40% at 40 DPI).

**Figure 3.**
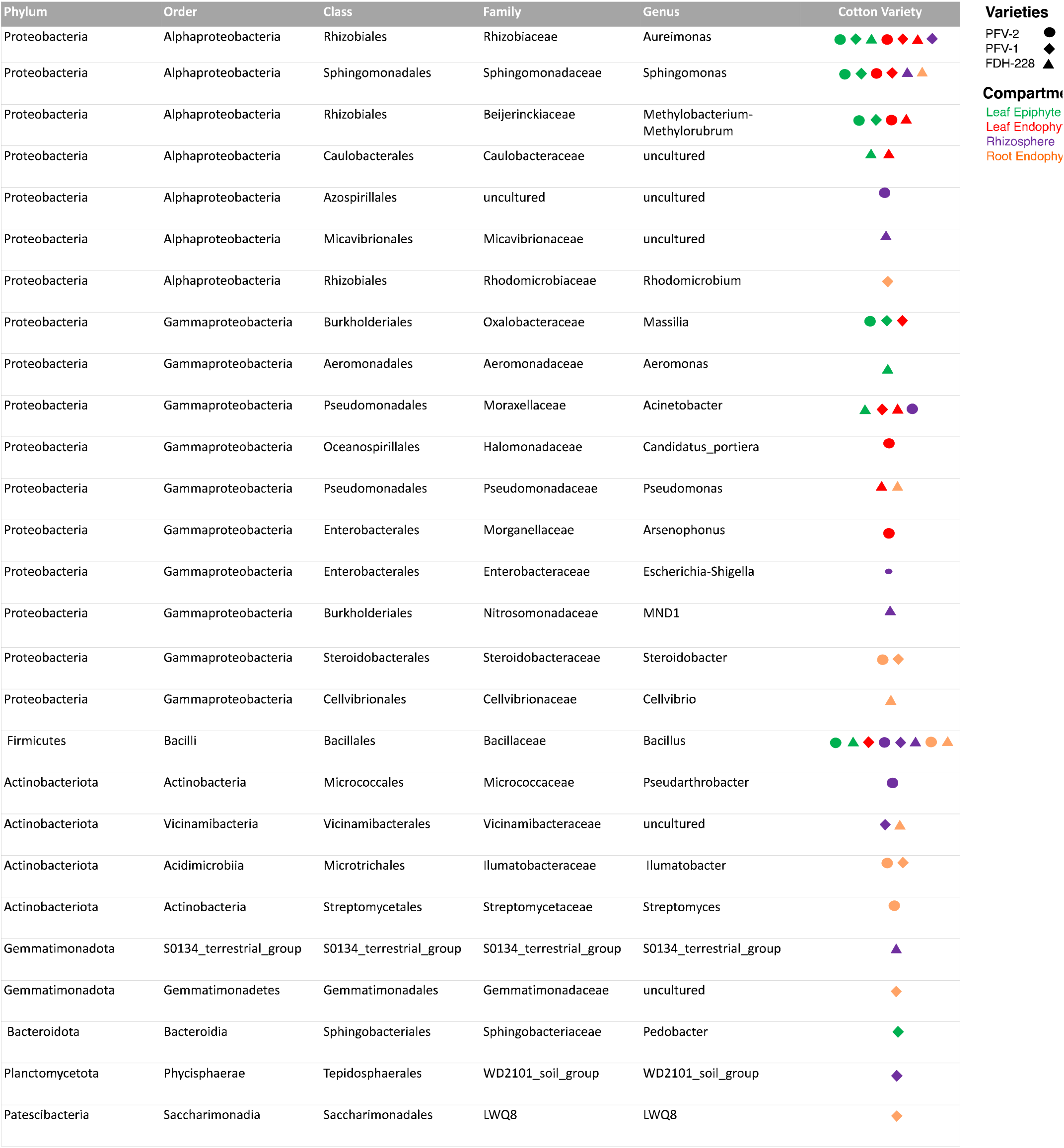
Top five most abundant core microbiome found in each plant Compartment of CLCuD susceptible, partially tolerant and tolerant cotton varieties. These are based on Supplementary Figures S2-S4 and represent the core microbiome at the right side of the heatmaps.

**Figure 4.**
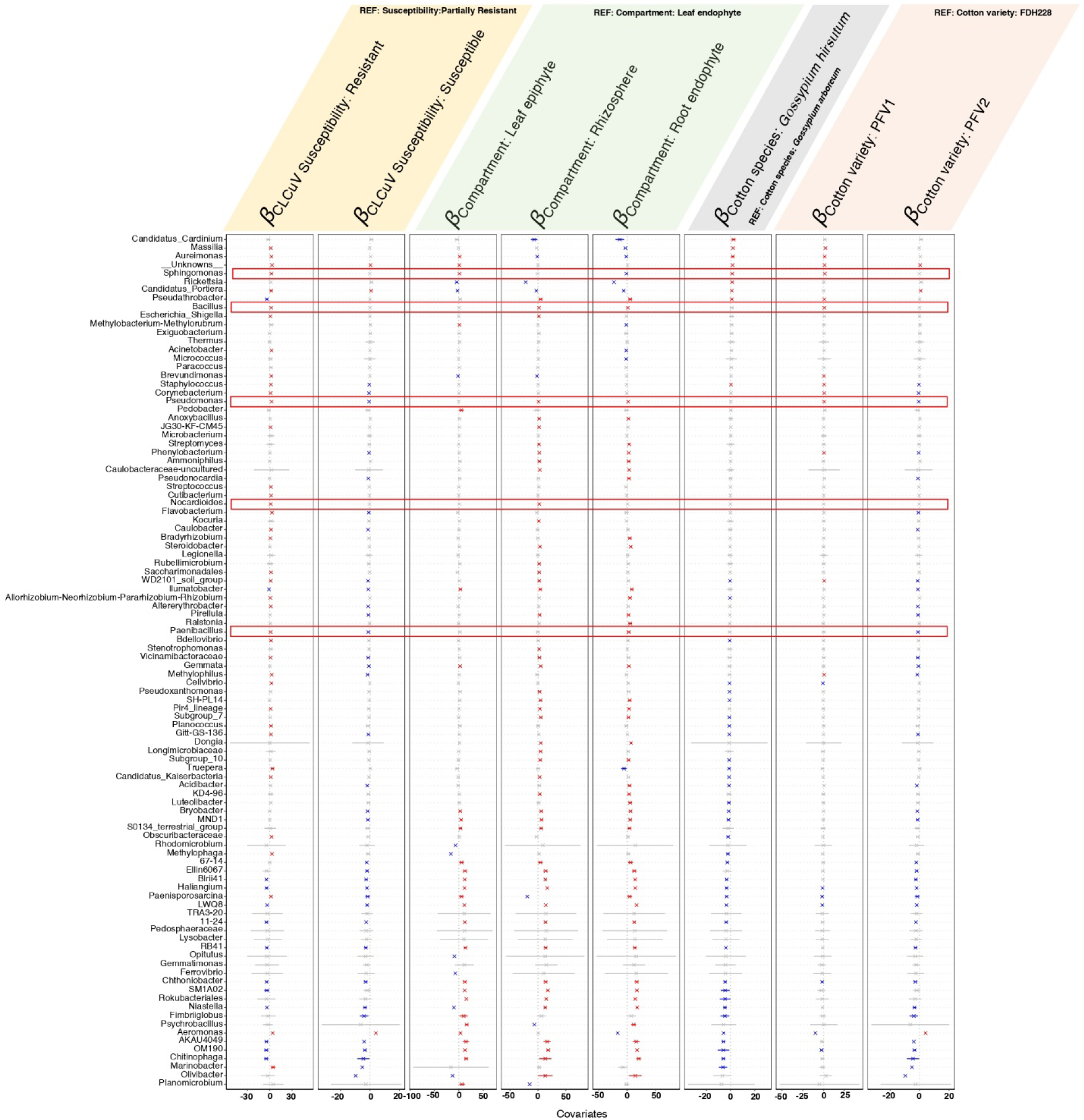
*β* -coefficients returned from GLLVM procedure for covariates considered in this comparative microbiome analysis between *Gossypium arboreum* and *Gossypium hirsutum* under CLCuD attack. Those coefficients which are positively associated with the microbial abundance of a particular species are represented in red colour whilst those that are negatively associated are represented with blue colour, respectively. Those microbes which were statistically insignificant, i.e., where the coefficients crossed 0 boundary, are greyed out. the Since the collation of ASVs were done at Genus level, all those ASVs that cannot be categorized based on taxonomy are collated under “ Unknowns “ category. The highlighted genera are salicylic acid producing bacteria.

**Figure 5.**
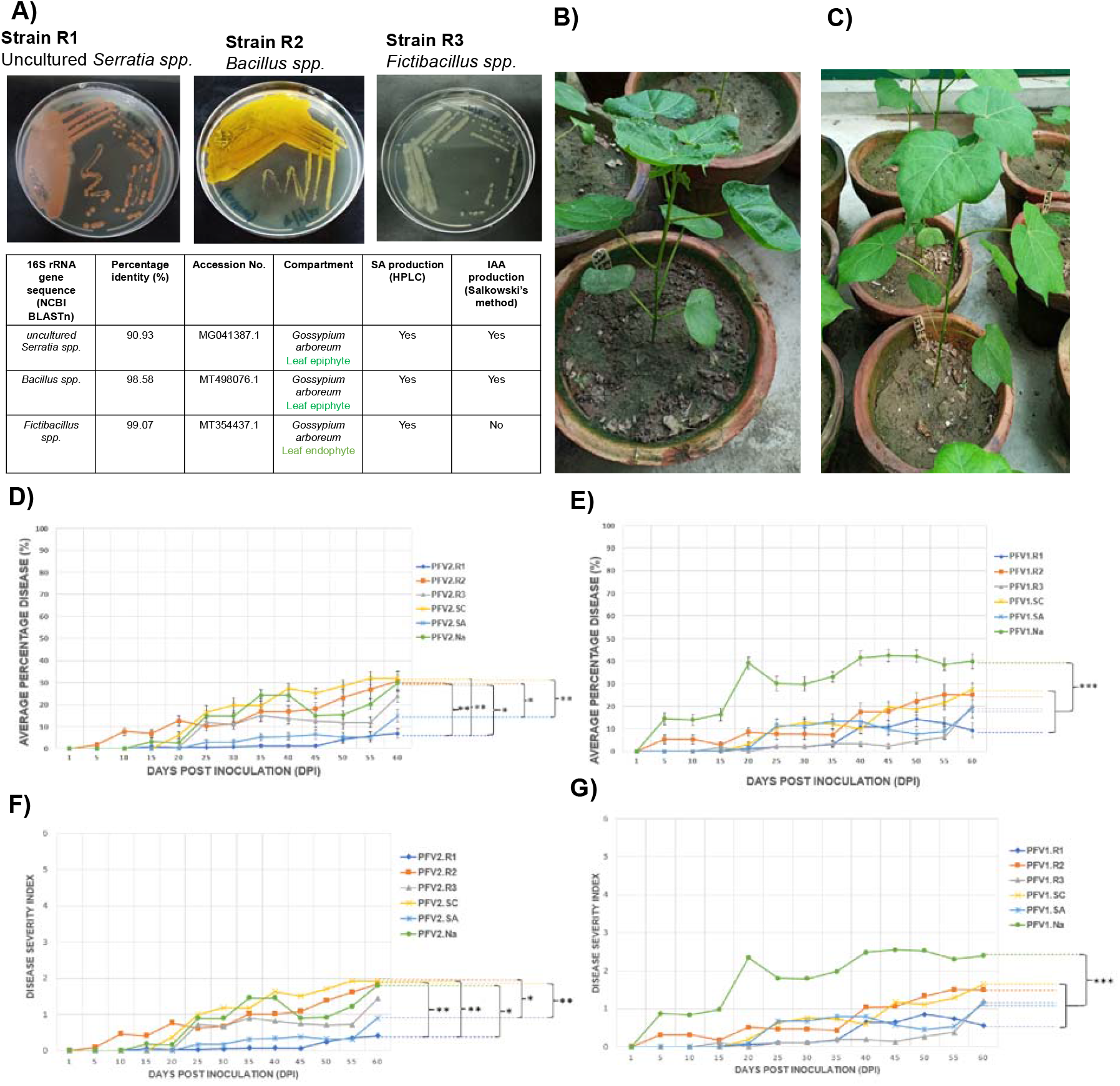
Microbiome mediated CLCuD suppression under SA producing strains Application. **A)** Salicylic Acid producing strains isolated from CLCuD tolerant variety FDH-228. **B)** *Gossypium hirsutum* PFV-1 control plant with no application (nA) exhibiting CLCuD symptoms at 20 DPI. **C)** *Gossypium hirsutum* PFV-1 plant with application of strain R1 (*uncultured Serratia spp*.) resisting disease at 20 DPI. **D)** CLCuD disease progression in Susceptible variety (PFV-2) under different applications: PFV2.R1 (*uncultured Serratia spp*.), PFV2.R2 (*Bacillus spp*.), PFV2.R3 (*Fictibacillus spp*.), PFV2.SC (Synthetic consortium of R1, R2, R3), PFV2.SA (Salicylic acid 400mg/L positive control), PFV2.NA (No application-control) Statistically significant levels are indicated by asterisks as * (p < 0.05), ** (p < 0.01) or *** (p < 0.001). **E)** CLCuD disease progression in Partially tolerant variety (PFV-1) under different applications: PFV1.R1 (*uncultured Serratia spp*.), PFV1.R2 (*Bacillus spp*.), PFV1.R3 (*Fictibacillus spp*.), PFV1.SC (Synthetic consortium of R1, R2, R3), PFV1.SA (Salicylic acid 400mg/L positive control), PFV1.NA (No application-control). **F)** and **G)** are similar to D) and E) except that we have represented the results in disease severity index. Statistically significant levels are indicated by asterisks as * (p < 0.05), ** (p < 0.01) or *** (p < 0.001).

## Discussion

CLCuD is an acute disease of cotton, with economic losses reaching approximately 2 billion USD per annum in Pakistan [31]. In comparison, the major losses happened in the 1990s, when cotton yields were reduced by 75% [32]. In the past five years, Pakistan has fallen from the 3rd largest [33] to the fifth largest cotton-producing nation [34], with CLCuD being the main contributor to this downfall. CLCuD is a whitefly-transmitted disease caused by the begomovirus. Pathogenic attacks constantly threaten plants, which pose a major global food security hazard. Over the past few years, extensive knowledge has been gathered from research on the effect of plant microbiome on fungal and bacterial pathogens. However, the role of the microbiome against viral pathogens remains unexplored. The endosphere and rhizosphere microbiome harbor potential microbes for suppressing plant diseases which can be exploited for combatting viral pathogens [35-37]. To our knowledge, this study is the first to investigate the phyllosphere and rhizospheric bacterial communities of CLCuV-infected susceptible, partially tolerant, and tolerant cotton plants. It is vital to understand the basis of resistance in the CLCuV-tolerant *G. arboreum* species, and here we present the microbiome angle of it.

The highest richness in taxonomy was observed in the RZ of the CLCuV-tolerant FDH-228. *Pseudomonas* was predominantly found in the RZ of CLCuV-tolerant *G. arboreum*. Rhizospheric microbiota in tomato plants is known to provide disease resistance against wilt caused by *Ralstonia solanacearum* [38]. Also, rhizobacteria belonging to the *Pseudomonadaceae, Bacillaceae, Solibacteraceae*, and *Cytophagaceae* families were reported to be prevalent in the rhizosphere of *Fusarium-*resistant bean cultivars [39]. No *bacillus* species were found in the leaf epiphytic region of CLCuV susceptible PFV-2. They are reported in the partially tolerant PFV-1 but most abundant in the tolerant variety FDH-228.

The temporal and spatial dynamics of microbial communities play a pivotal role in understanding the underlying mechanisms of plant defense. Induced systemic resistance (ISR) is also activated in plants in an SA-dependent manner. *Pseudomonas aeruginosa* 7NSK2 failed to activate ISR in SA mutant tomato plants [40]. In the core microbiome analysis, *Pseudomonas* has been found in the LN and RN of *G. arboreum*. Endophytic bacteria such as *Bacillus, Pseudomonas*, and *Rhizobium* have been reported to confer pathogen resistance by the production of bioactive compounds [41-47]. *Pseudomonas*, which has been observed to be a key player in the CLCuD-tolerant cotton variety’s microbiome, is known to synthesize SA from chorismate via the isochorismate synthase (ICS) and isochorismate pyruvate lyase [48, 49]. *Pseudomonas* and *Bacillus* are known to be positively associated with CLCuV resistance, as exhibited in our GLLVM results. Other bacterial species known for endogenous SA production and enhancing plant host defenses by activation of SAR include *P. fluorescens* (strain CHA0), *P. aeruginosa* (7NSK2), and *Serratia marcescens* (strain 90-166) [50–55].

The three identified SA-producing bacterial strains have been isolated from the CLCuD tolerant variety FDH-228. The application of bacterial strains from the resistant variety aids the susceptible *Gossypium hirsutum* in resisting disease compared to control plants. When SA levels increase after pathogen infection during basal resistance, SA binds to NPR4 and releases more NPR1, which activates the SAR. At an extremely high SA level during the hypersensitive response (HR), SA binds to NPR3 and promotes its interaction with NPR1, which finally leads to the turnover of NPR1 [56, 57]. Salicylic acid (SA) is synthesized after inoculating plants with pathogens or exposure to certain abiotic stresses, such as ozone and UV-C light. SA was found to be essential for gene-for-gene resistance, systemic acquired resistance, and reduction of disease development after inoculation with virulent pathogens [58]. In a study conducted by [59], the rhizobacterial strain *Serratia marcescens* 90-166 mediated induced systemic resistance (ISR) to fungal, bacterial, and viral pathogens.

Endophytic bacteria exhibit the potential to trigger salicylic acid-dependent resistance and alter the rhizospheric bacterial communities. In another study [60], the possible roles of endophytic *Bacillus megaterium* in rice resistance against rice spikelet rot disease (SRD) were elucidated, and inferences were made on the potential interactions between plant defenses and the rhizosphere microenvironment. A correlation between plant resistance and the rhizosphere microbiome was observed.

The plant-pathogen and plant-microbe interactions represent how a plant immune system would react to a pathogen attack. Therefore, it is mandatory to explore the microbiome itself and the ‘microbiome under pathogenic attack,’ which reveals the answers to the overall success or failure of the plant-beneficial microbiome. Microbiota use may have a probiotic effect on plant functions, as the basis of plant defense is also a function of the plant genome and its microbial counterpart. Using the plant’s microbiome for suppressing begomoviral disease seems very promising. The SA-producing bacterial isolates were applied as a biocontrol agent, strain R1 (uncultured *Serratia sp*.) was performed, and the positive control group where SA was applied as a protective regime in the susceptible variety PFV-2. The average percentage of disease in partially tolerant variety PFV-1 remained less than 20% till 60 days post-viral inoculation (DPI) in plants applied with strain R1 (uncultured *Serratia sp*.), strain R3 (*Fictibacillus spp*.), and SA respectively as compared to control (no application) with 40% disease progression at 60 DPI. Thus, the CLCuD-resistant variety-associated microbiome has the profound potential to resist disease under high whitefly attack in the highly susceptible *Gossypium hirsutum*.

## Conclusion

High Salicylic Acid (SA) production in bacterial strains isolated from the phyllosphere of Cotton Leaf Curl Disease (CLCuD) tolerant *Gossypium arboreum* exhibited the ability of these strains to activate the plant defense responses prior to viral pathogenic attack. Compared to the control, these strains proved efficient in suppressing viral disease in CLCuD-susceptible cotton species. The plants applied with *Serratia* and *Fictibacillus* strains in monoassociation experiments proved the most effective in CLCuD suppression. The microbiome analysis of *Gossypium hirsutum* and *Gossypium arboreum* has provided us with insights into the potential of the microbiome associated with the tolerant species, and the biocontrol agents identified in this study have the potential to be utilized in intervention studies for field applications against CLCuD.

## Author Contributions

Rhea Aqueel (Conceptualization, Visualization, Data Curation, Investigation, Formal analysis, Writing - Original Draft)

Ayesha Badar (Data Curation, Visualization, Formal analysis, Investigation, Writing - Review & Editing)

Nazish Roy (Conceptualization, Methodology, Writing - Review & Editing, Supervision, Funding Acquisition)

Qandeel Mushtaq (Investigation, Writing - Review & Editing)

Aimen Fatima Ali (Investigation, Writing - Review & Editing)

Aftab Bashir (Resources, Writing - Review & Editing)

Umer Zeeshan Ijaz (Methodology, Software, Formal analysis, Writing - Original Draft, Supervision, Funding Acquisition)

Kauser Abdulla Malik (Conceptualization, Methodology, Writing – review & editing, Project Administration, Supervision, Funding Acquisition)

## Acknowledgments

We would like to thank all the field and laboratory teams of Kauser Abdullah Malik School of Life Sciences and Dr Rashid Masih (Chemistry Department, FCCU) for his contributions towards helping in HPLC. We are grateful to Four Brothers Research Group, Pakistan and Ayub Agricultural Research Institute (AARI), Faisalabad for providing the cotton seeds for our pot experiment analysis.

## Data Availability

All the sequencing data and relevant meta data is available from the corresponding authors on reasonable request.

## References

[1] Rahman, M. U., Khan, A. Q., Rahmat, Z., Iqbal, M. A., & Zafar, Y. (2017). Genetics and genomics of cotton leaf curl disease, its viral causal agents and whitefly vector: a way forward to sustain cotton fiber security. Frontiers in Plant Science, 8, 1157.

[2] Mansoor, S., Briddon, R. W., Bull, S. E., Bedford, I. D., Bashir, A., Hussain, M., Saeed, M., Zafar, Y., Malik, K.A., Fauquet, C. & Markham, P. G. (2003). Cotton leaf curl disease is associated with multiple monopartite begomoviruses supported by single DNA β. Archives of virology, 148(10), 1969–1986.

[3] Razaq M, Abbas G, Farooq M, Aslam M, Athar H-u-R. Effect of insecticidal application on aphid population, photosynthetic parameters and yield components of late sown varieties of canola, Brassica napus L. Pakistan Journal of Zoology. 2014; 46(3):661–8. 5.

[4] Oerke E-C. Crop losses to pests. The Journal of Agricultural Science. 2006; 144(01):31–43. 4.

[5] Rahman, M. Zafar Y (2007a) Registration of NIBGE-115. J Plant Regist, 1, 51–52.

[6] Wendel, J. F., Brubaker, C., Alvarez, I., Cronn, R., & Stewart, J. M. (2009). Evolution and natural history of the cotton genus. In Genetics and genomics of cotton (pp. 3–22). Springer, New York, NY.

[7] Tlais, A. Z. A., Lemos Junior, W. J. F., Filannino, P., Campanaro, S., Gobbetti, M., & Di Cagno, R. (2022). How microbiome composition correlates with biochemical changes during sauerkraut fermentation: A focus on neglected bacterial players and functionalities. Microbiology spectrum, 10(4), e00168–22.

[8] Roy N, Choi K, Khan R, Lee SW (2019). Culturing Simpler and Bacterial Wilt Suppressive Microbial Communities from Tomato Rhizosphere. The plant pathology journal; 35(4):362–371.

[9] Wei, F., Zhao, L., Xu, X., Feng, H., Shi, Y., Deakin, G., … & Zhu, H. (2019). Cultivar-dependent variation of the cotton rhizosphere and endosphere microbiome under field conditions. Frontiers in Plant Science, 1659.

[10] Turner, T. R., James, E. K., & Poole, P. S. (2013). The plant microbiome. Genome biology, 14(6), 1–10.

[11] Rossmann, M., Sarango-Flores, S.W., Chiaramonte, J.B., Kmit, M.C.P., Mendes, R., 2017. Plant microbiome: Composition and functions in plant compartments. In: Pylro, V., Roesch, L. (Eds.), The Brazilian Microbiome, Springer, Cham, pp. 7–20.

[12] Grube, M., Cernava, T., Soh, J., Fuchs, S., Aschenbrenner, I., Lassek, C., Wegner, U., Becher, D., Riedel, K., Sensen, C.W., Berg, G., 2015. Exploring functional contexts of symbiotic sustain within lichen-associated bacteria by comparative omics. ISME J. 9, 412–424.

[13] Qiao, Q., Wang, F., Zhang, J., Chen, Y., Zhang, C., Liu, G., Zhang, H., Ma, C & Zhang, J. (2017). The variation in the rhizosphere microbiome of cotton with soil type, genotype and developmental stage. Scientific reports, 7(1), 1–10.

[14] Zhang, J., Liu, Y.-X., Zhang, N., Hu, B., Jin, T., Xu, H., et al. (2019). NRT1. 1B is associated with root microbiota composition and nitrogen use in field-grown rice. Nat. Biotechnol. 37, 676–684. doi: 10.1038/s41587-019-0104-4

[15] Stringlis, I. A., Yu, K., Feussner, K., De Jonge, R., Van Bentum, S., Van Verk, M. C., et al. (2018). MYB72-dependent coumarin exudation shapes root microbiome assembly to promote plant health. Proc. Natl. Acad. Sci. 115, E5213–E5222. doi: 10.1094/PBIOMES-11-18-0050-R

[16] Yim, W., Seshadri, S., Kim, K., Lee, G., Sa, T.M., 2013. Ethylene emission and PR protein synthesis in ACC deaminase producing Methylobacterium spp. inoculated tomato plants (Lycopersicon esculentum Mill.). Plant Physiol. Biochem. 67, 95–104.

[17] Agafonova, N.V., Doronina, N.V., Trotsenko, Y.A., 2016. Enhanced resistance of pea plants to oxidative stress caused by paraquat during colonization by aerobic Methylobacteria. Appl. Biochem. Microbiol. 52 (2), 199–204.

[18] Mishra, S., Singh, A., Keswani, C., Saxena, A., Sarma, B.K., Singh, H.B., 2015. Harnessing plant-microbe interactions for enhanced protection against phytopathogens. In: Arora, N.K. (Ed.), Plant Microbes Symbiosis: Applied Facets, Springer, New Delhi, pp. 111–125.

[19] Ardanov, P., Sessitsch, A., Haggman, H., Kozyrovska, N., Pirttila, A.M., 2012. Methylobacterium-induced endophyte ommunity changes correspond with protection of plants against pathogen attack. PLoS One. 7(10)e46802

[20] Bulgarelli, D., Schlaeppi, K., Spaepen, S., Van Themaat, E. V. L., & Schulze-Lefert, P. (2013). Structure and functions of the bacterial microbiota of plants. Annual review of plant biology, 64, 807–838.

[21] Guttman, D. S., McHardy, A. C., & Schulze-Lefert, P. (2014). Microbial genome-enabled insights into plant–microorganism interactions. Nature Reviews Genetics, 15(12), 797–813.

[22] Wagi, S., & Ahmed, A. (2017). Phyllospheric plant growth promoting bacteria. J Bacteriol Mycol, 5, 215–216.

[23] Archana, K., Kaur, S. M., Pashupat, V., Javed, A., & Dharmender, P. (2020). Role of 2, 6 Dichloroisonicotinic acid inducing resistance in cotton against cotton leaf curl disease. Research Journal of Biotechnology Vol, 15, 5.

[24] Howe, G. A., & Jander, G. (2008). Plant immunity to insect herbivores. Annu. Rev. Plant Biol., 59, 41–66.

[25] Carvalho, T. L. G., Balsemão-Pires, E., Saraiva, R. M., Ferreira, P. C. G., & Hemerly, A. S. (2014). Nitrogen signalling in plant interactions with associative and endophytic diazotrophic bacteria. Journal of experimental botany, 65(19), 5631–5642.

[26] Rossmann, M., Sarango-Flores, S. W., Chiaramonte, J. B., Kmit, M. C. P., & Mendes, R. (2017). Plant microbiome: composition and functions in plant compartments. In The Brazilian Microbiome (pp. 7–20). Springer, Cham.

[27] Zheng, W., Tsompana, M., Ruscitto, A., Sharma, A., Genco, R., Sun, Y., & Buck, M. J. (2015). An accurate and efficient experimental approach for characterization of the complex oral microbiota. Microbiome, 3(1), 48.

[28] Ehmann, A.K. (1977). The van urk-Salkowski reagent--a sensitive and specific chromogenic reagent for silica gel thin-layer chromatographic detection and identification of indole derivatives. Journal of chromatography, 132 2, 267–76.

[29] Chen, C., Bélanger, R. R., Benhamou, N., & Paulitz, T. C. (1999). Role of salicylic acid in systemic resistance induced by Pseudomonas spp. against Pythium aphanidermatum in cucumber roots. European Journal of Plant Pathology, 105, 477–486.

[30] Archana, K., Kaur, S. M., Pashupat, V., Javed, A., & Dharmender, P. (2020). Role of 2, 6 Dichloroisonicotinic acid inducing resistance in cotton against cotton leaf curl disease. Research Journal of Biotechnology Vol, 15, 5.

[31] Rafiq, A., Ali, W. R., Asif, M., Ahmed, N., Khan, W. S., Mansoor, S., … & Amin, I. (2021). Development of a LAMP assay using a portable device for the realtime detection of cotton leaf curl disease in field conditions. Biology Methods and Protocols, 6(1), bpab010.

[32] Mansoor, S., Amin, I., & Briddon, R. W. (2011). Geminiviral diseases of cotton. StreSS PhySiology in Cotton, 125.

[33] [Rana, A. W., Ejaz, A., & Shikoh, S. H. (2020). Cotton crop: A situational analysis of Pakistan. Intl Food Policy Res Inst.

[34] SOHAİL, N., Sarfaraz, Y., & RafİQue, A. (2023). PAKISTAN FLOODS: AN INSIGHT INTO AGRICULTURE AND FOOD SUPPLY. Trakya University Journal of Natural Sciences, (Online First Publications)].

[35] Mendes, R.,,, Kruijt, M.,,, De Bruijn, I.,,, Dekkers, E.,,, Van Der Voort, M.,,, Schneider, J.H.,,, Piceno, Y.M.,,, DeSantis, T.Z.,,, Andersen, G.L.,,, Bakker, P.A.,,, et al. Deciphering the rhizosphere microbiome for disease-suppressive bacteria. Science 2011, 332, 1097–1100. [CrossRef]

[36] Durán, P.,,, Jorquera, M.,,, Viscardi, S.,,, Carrion, V.J.,,, de la Luz Mora, M.,,, Pozo, M.J. Screening and characterization of potentially suppressive soils against gaeumannomyces graminis under extensive wheat cropping by chilean indigenous communities. Front. Microbiol. 2017, 8, 1552. [CrossRef]

[37] Carrión, V.J.,,, Cordovez, V.,,, Tyc, O.,,, Etalo, D.W.,,, de Bruijn, I.,,, de Jager, V.C.,,, Medema, M.H.,,, Eberl, L.,,, Raaijmakers, J.M. Involvement of Burkholderiaceae and sulfurous volatiles in disease-suppressive soils. ISME J. 2018, 12, 2307–2321. [CrossRef]

[38] Kwak, M.J.,,, Kong, H.G.,,, Choi, K.,,, Kwon, S.K.,,, Song, J.Y.,,, Lee, J.,,, Lee, P.A.,,, Choi, S.Y.,,, Seo, M.,,, Lee, H.J.,,, et al. Rhizosphere microbiome structure alters to enable wilt resistance in tomato. Nat. Biotechnol. 2018, 36, 1100–1109. [CrossRef] [PubMed]

[39] Mendes, L.W.,,, Raaijmakers, J.M.,,, De Hollander, M.,,, Mendes, R.,,, Tsai, S.M. Influence of resistance breeding in common bean on rhizosphere microbiome composition and function. ISME J. 2018, 12, 212–224. [CrossRef] [PubMed]

[40] McManus, P.S.,,, Stockwell, V.O.,,, Sundin, G.W.,,, Jones, A.L. Antibiotic use in plant agriculture. Annu. Rev. Phytopathol. 2002, 40, 443–465. [CrossRef]

[41] Meguro, A.,,, Ohmura, Y.,,, Hasegawa, S.,,, Shimizu, M.,,, Nishimura, T.,,, Kunoh, H. An endophytic actinomycete, Streptomyces sp. MBR-52, that accelerates emergence and elongation of plant adventitious roots. Actinomycetologica 2006, 20, 1–9. [CrossRef]

[42] Gibert, A.,,, Volaire, F.,,, Barre, P.,,, Hazard, L. A fungal endophyte reinforces population adaptive differentiation in its host grass species. New Phytol. 2012, 194, 561–571. [CrossRef] [PubMed

[43] Shimizu, M.,,, Nakagawa, Y.,,, Sato, Y.,,, Furumai, T.,,, Igarashi, Y.,,, Onaka, H.,,, Yoshida, R.,,, Kunoh, H. Studies on Endophytic Actinomycetes (I) Streptomyces sp. isolated from Rhododendron and Its antifungal activity. J. Gen. Plant Pathol. 2000, 66, 360–366. [CrossRef]

[44] Rodriguez, R.,,, Redman, R. More than 400 million years of evolution and some plants still can’t make it on their own: Plant stress tolerance via fungal symbiosis. J. Exp. Bot. 2008, 59, 1109–1114. [CrossRef]

[45] Conn, V.M.,,, Walker, A.R.,,, Franco, C.M.M. Endophytic actinobacteria induce defense pathway in Arab. thaliana. Mol. Plant Microbe Interact. 2008, 21, 208–218. [CrossRef]

[46] Khalaf, E.M.,,, Raizada, M.N. Bacterial seed endophytes of domesticated cucurbits antagonize fungal and oomycete pathogens including powdery mildew. Front. Microbiol. 2018, 9, 1–18.

[47] Terhonen, E.,,, Blumenstein, K.,,, Kovalchuk, A.,,, Asiegbu, F.O. Forest tree microbiomes and associated fungal endophytes: Functional roles and impact on forest health. Forests 2019, 10, 1–33. [CrossRef]

[48] Wildermuth, M.C., Dewdney, J., Wu, G. and Ausubel, F.M., 2001. Isochorismate synthase is required to synthesize salicylic acid for plant defence. Nature, 414(6863), pp.562–565.

[49] Serino, L.,,, Reimmann, C.,,, Baur, H.,,, Beyeler, M.,,, Visca, P.,,, Haas, D. Structural genes for salicylate biosynthesis from chorismate in Pseudomonas aeruginosa. Mol. Gen. Genet. 1995, 249, 217–228. [CrossRef]

[50] De Meyer, G.,,, Höfte, M. Salicylic acid produced by the rhizobacterium Pseudomonas aeruginosa 7NSK2 induces resistance to leaf infection by Botrytis cinerea on bean. Phytopathology 1997, 87, 588–593. [CrossRef]

[51] Press, C.M.,,, Wilson, M.,,, Tuzun, S.,,, Kloepper, J.W. Salicylic acid produced by Serratia marcescens 90-166 is not the primary determinant of induced systemic resistance in Cucumber or Tobacco. Mol. Plant Microbe Interact. 1997, 10, 761–768. [CrossRef]

[52] An, C.,,, Mou, Z. Salicylic acid and its function in plant immunity. J. Integr. Plant Biol. 2011, 53, 412–428. [CrossRef] [PubMed] Microorganisms 2020, 8, 31 13 of 15

[53] Anand, A.,,, Uppalapati, S.R.,,, Ryu, C.M.,,, Allen, S.N.,,, Kang, L.,,, Tang, Y.,,, Mysore, K.S. Salicylic acid and systemic acquired resistance play a role in attenuating crown gall disease caused by Agrobacterium tumefaciens. Plant Physiol. 2008, 146, 703–715. [CrossRef] [PubMed]

[54] Denancé, N.,,, Sánchez-Vallet, A.,,, Goffner, D.,,, Molina, A. Disease resistance or growth: The role of plant hormones in balancing immune responses and fitness costs. Front. Plant Sci. 2013, 4, 1–12. [CrossRef] [PubMed]

[55] Yang, D.L.,,, Yang, Y.,,, He, Z. Roles of plant hormones and their interplay in rice immunity. Mol. Plant 2013, 6, 675–685. [CrossRef]

[56] Fu, Z. Q., Yan, S., Saleh, A., Wang, W., Ruble, J., Oka, N., … & Dong, X. (2012). NPR3 and NPR4 are receptors for the immune signal salicylic acid in plants. Nature, 486(7402), 228–232.

[57] Moreau, M., Tian, M., & Klessig, D. F. (2012). Salicylic acid binds NPR3 and NPR4 to regulate NPR1-dependent defense responses. Cell research, 22(12), 1631–1633.

[58] Nawrath, C., Heck, S., Parinthawong, N., & Métraux, J. P. (2002). EDS5, an essential component of salicylic acid–dependent signaling for disease resistance in Arabidopsis, is a member of the MATE transporter family. The Plant Cell, 14(1), 275–286.

[59] Press, C. M., Wilson, M., Tuzun, S., & Kloepper, J. W. (1997). Salicylic acid produced by Serratia marcescens 90-166 is not the primary determinant of induced systemic resistance in cucumber or tobacco. Molecular Plant-Microbe Interactions, 10(6), 761–768.

[60] Cheng, T., Yao, X. Z., Wu, C. Y., Zhang, W., He, W., & Dai, C. C. (2020). Endophytic Bacillus megaterium triggers salicylic acid-dependent resistance and improves the rhizosphere bacterial community to mitigate rice spikelet rot disease. Applied Soil Ecology, 156, 103710.

