## Supplemental information for "Cotton Microbiome Profiling and Cotton Leaf Curl Disease (CLCuD) Suppression through Microbial Consortia associated with *Gossypium arboreum*"

**e-Supplementary Methods**

**Beta Satellite Amplification for CLCuV confirmation**

Beta satellite DNA component of the CLCuD complex was amplified using the primers listed in Supplementary Table S1. For each reaction, 25 µL total volume was used with 12.5 µL Thermo Scientific DreamTaq Green PCR Master Mix (2X), 1 µL from 10 µM of each primer, 1 µL of $MgCl_2$, 1 µL of total genomic DNA template (10 ng/µL), and remaining volume was made up with nuclease free water. The PCR program was set as: 94 ˚C for 4 min (initial denaturation), followed by 30 cycles of 94 ˚C for 1 min (denaturation), 68 ˚C for 1 min (annealing), 72 ˚C for 1 min 30 sec (extension) with a final extension of 10 min at 72 ˚C.

**Bioinformatics Analysis**

Abundance tables were generated by constructing amplicon sequencing variants (ASVs) using the Qiime2 workflow with Deblur algorithm (1) after consulting https://qiita.ucsd.edu/static/doc/html/deblur_quality.html for recommendations. Briefly, the sequencing reads were imported to Qiime2 format, and were quality trimmed with a minimum Phred quality score of 20. Afterwards, we have used the qiime deblur denoise-other plugin with parameters --p-trim-length 280 --p-min-size 2 --p-min-reads 2 to generate ASVs. As a pre-processing step, the method also filters out any sequences that are not found in the reference

SILVA SSU Ref NR database v138 which, is additionally used in qiime feature-classifier plugin to assign taxonomy to each ASV. This yielded a 59 (sample) X 38,120 (ASV) abundance table with summary statistics of sample reads as follows: [1st Quartile:7,979; Median:15,522; Mean: 14;565; 3rd Quartile:21,387; and Maximum: 27,839]. Afterwards qiime phylogeny align-to-tree-mafft-fasttree generated the rooted phylogenetic tree of all the ASVs. The biom file for the ASVs was generated by combining the abundance table with taxonomy information using biom utility available in qiime2 workflow. Next we used PICRUSt2 (2) and its qiime2 plugin [https://github.com/gavinmdouglas/q2-picrust2] using the parameters --p-hsp-method pic --p-max-nsti 2 in qiime picrust2 full-pipeline to find KEGG enzymes and MetaCyc pathway predictions. For ensuing statistical analysis, any sample having <5000 total reads was dropped, excluding further contaminants based on taxonomy (Chloroplast, Mitochondria, and ASVs unassigned at Phylum level).

**Statistical Analysis**

Statistical analyses were performed in R using the combined data generated from the bioinformatics as well as meta data associated with the study. The R's Vegan package (3) was used for the analysis of alpha diversity of all tables. For alpha diversity, the indices used were (i) rarefied richness – the number of expected features in a rarefied sample (to the minimum library size, and (ii) Shannon entropy – an index that takes into account both richness and diversity to provide a measurement of community balance. For beta diversity, the dissimilarity in species community composition between pairwise comparisons of bacterial communities were represented in Principal Coordinate Analysis (PCoA) ordination plots by using three different distance metrices in Vegan 's cmdscale function: (i) Bray Curtis, which considers the species abundance count; (ii) Unweighted Unifrac, which considers the phylogenetic distance between the branch lengths of ASVs observed in different samples (implemented in the phyloseq package (4); and (iii) Hierarchical Meta-Storms (HMS), (5) a functional beta diversity distance which collapses the metabolic pathways at the observed KOs level by considering BRITE pathways as a set of reference pathways, and then propagating the abundances upward for these pathways in a multi-level pathway hierarchy to give a weighted dissimilarity measure. To see if beta diversity is statistically significant between the groups, we have used Vegan's adonis for analysis of variance using distance matrices (BrayCurtis/Unweighted Unifrac/Hierarchical Meta-Storms) This function, henceforth referred to as PERMANOVA, fits linear models to distance matrices and uses a permutation test with pseudo-F ratios and gives an $R^2$ value for each covariate defined as percentage variability in community structure explained by that covariate.

To find the core microbiome, we have used R's microbiome package (6) which offers a two-dimensional representation of core communities by taking an assumed initial prevalence (50% presence of a microbe in the total number of samples) and then calculating the detection limit in terms of abundances to differentiate between low-abundance core microbiome versus high-abundance core microbiome. The initial prevalence threshold was set to 50%. In addition, we have generated taxa bar plots of the top 20 most abundant phyla at appropriate taxonomic levels to give an indication of how dominant phyla change between conditions.

To see the relationship between different conditions (cotton plant varieties and compartments) and the minimal subset of microbes that can explain them, we used the variable selection approach where through penalized regression on the set of all pairwise log-ratios (35) we identify two disjoint subsets of microbes, those that are positively associated, and those that are negatively associated with the covariate of interest. Briefly, we used the CODA-LASSO approach (7) where the abundance of individual covariate $y_i$ is modeled as $y_i = \beta_0 + \beta_1 \log(x_{1i}) + \cdots + \beta_j \log(x_{ji}) + \epsilon_i$ (for $i$-th sample and $j$-th species/function, with $x_{ji}$ being the microbe abundance) with the constraint $\sum_{k\geq 1} \beta_k = 0$ (i.e., all $\beta$-coefficients sum up to 1), and these regression coefficients $\boldsymbol{\beta} = (\beta_0, \ldots, \beta_j)$ are estimated to minimise $\sum_{i=1}^{n}(y_i - \beta_0 - \beta_1 \log(x_{1i}) - \cdots - \beta_j \log(x_{ji}))^2 + \lambda \sum_{k\geq 1}|\beta_k|$ subject to $\sum_{k\geq 1} \beta_k = 0$ (using a soft thresholding and projection algorithm) for $n$ samples. Here, $\lambda$ is the penalization parameter in Lasso shrinkage terms $\lambda \sum_{k\geq 1}|\beta_k|$ which forces some of the $\boldsymbol{\beta}$-coefficients to go zero, particularly those that do not have a relationship with the covariates and serves as a means to do variable selection. The non-zero $\boldsymbol{\beta}$-coefficients are then divided into two groups, those that are positively associated with the environmental covariate, and those that are negatively associated with the environmental covariate, respectively. For this purpose, we used coda glmnet() function from R's coda4microbiome package (8). We have used the top 100 most abundant genera/functions in the CODA-LASSO model. For visualizing the expression of microbes/functions selected from the procedure, we have used TSS+CLR normalization (Total Sum Scaling followed by Centralized Log Ratio).

To find the relationship between microbial communities and sources of variation (Cotton Species, Cotton Variety, CLCuV_Susceptibility, and Compartment), we have used Generalised Linear Latent Variable Model (GLLVM) (9) which extends the basic generalized linear model that regresses the mean abundances $\mu_{ij}$ (for $i$-th sample and $j$-th microbe) of individual microbes against environmental covariates $x_i$ as above by incorporating latent variables $u_i$ as $g(\mu_{ij}) = \eta_{ij} = \alpha_i + \beta_{0j} + \boldsymbol{x}_i^T \boldsymbol{\beta}_j + \boldsymbol{u}_i^T \boldsymbol{\theta}_j$, where $\boldsymbol{\beta}_j$ are the microbe specific coefficients associated with individual covariate (a 95% confidence interval of these whether positive or negative, and not crossing 0 boundary gives directionality with the interpretation that an increase or decrease in that particular covariate causes an increase or decrease in the abundance of the microbe), and $\boldsymbol{\theta}_j$ are the corresponding coefficients associated with latent variable. $\beta_{0j}$ are microbes' specific intercepts, whilst $\alpha_i$ are optional sample effects which can either be chosen as fixed effects or random effects. To model the distribution of individual microbes, we have used Negative Binomial distribution. Additionally, the approximation to the log-likelihood is done through Laplace approximation (LA) with final sets of parameters in glvmm() function being family = 'negative.binomial', method="LA", and starting.val='zero' that seemed to fit well. This, we did for top 100 most abundant genera in our datasets. In addition, the factor loadings $\boldsymbol{\theta}_j$ store correlations of microbes with the residual covariance matrix $\boldsymbol{\Sigma} = \boldsymbol{\Gamma}\boldsymbol{\Gamma}^T$ where $\boldsymbol{\Gamma} = [\theta_1 \dots \theta_m]$ for $m$ latent variables. This residual covariance matrix gave co-occurrence relationship between microbes that are not explained by environmental covariates as above.

All figures in this study were generated using R's ggplot2 package (10). For alpha diversity we have used ANOVA, and where two categories are significantly different, following annotations are used to denote significance: '***' ($p \leq 0.001$), '**' ($p \leq 0.01$), '*' ($p \leq 0.05$), and '.' ($p \leq 0.1$).

**Morphological & Biochemical Characterization of Isolated Bacteria**

Cell morphology was studied under light microscope. Gram staining (11) was used to differentiate between Gram-positive and Gram-negative bacteria. Motility of bacterial isolates was observed under light microscope. Sterilized distilled water was used to prepare bacterial suspension. A drop of suspension was added to the glass slide and observed under the microscope.

Quick test strips (QTS) 24 bacterial identification kits (DESTO Laboratories, Karachi Pakistan) were used to detect enzymes and carbon source utilization pattern. Bacterial suspension was prepared in sterilized falcon tubes from freshly streaked colonies. Incubation box of QTS strips were partially filled with water to make the inside environment of QTS-24 strips moist. QTS-24 strips were placed in the incubation boxes. All the cups were partially filled with except cups of UREA and MOT (Cell Motility) that were completely filled with bacterial suspension. A gelatin disc was placed in cup labeled with GEL in each strip. Sterilized liquid paraffin (Mineral oil) was poured in cup labeled with ADH (Arginine deaminases) and $H_2S$ for an-aerobiosis. Inoculated QTS-24 strips were incubated at 28° C. After 24 h incubation, strips were examined, and the results were interpreted according to the manufacturer's manual. For TDA (Tryptophan deaminase) test, 1-2 drops of 10% ferric chloride 12 was added in TDA labeled cup. 0.8% sulphonic acid and 0.5% alpha-naphthylamine in 5N acetic acid was used for the detection of nitrate reductase test. For VP test, 40% KOH and 5% ethanolic solution of alpha-naphthol was added in VP labeled cup and incubated at 37 ° C for 10 min. For indole test, single drop of Kovac's reagent was added into cup labeled as IND (indole-3-acetic acid).

**Supplementary Table S1.** Primers used in the study

| Target Gene | Primer Sequence | Amplicon Size | Reference |
|---|---|---|---|
| **16S rRNA V3-V4 with Illumina adapter hang sequences** | 341F (5′-TCGTCGGCAGCGTCAGATGTGTATAAGAGACAGCCTACGGGNGGCWGCAG-3′)<br>805R (5′-GTCTCGTGGGCTCGGAGATGTGTATAAGAGACAGGACTACHVGGGTATCTAATCC-3′) | ~467 bp | (12) |
| **16S rRNA V3-V4 for Sanger Sequencing** | 341F (5'- CCTACGGGNGGCWGCAG -3')<br>805R (5'- GACTACHVGGGTATCTAATCC -3') | ~444bp | (12) |
| **Beta-Satellite DNA** | F-5'- GGTACCACTACGCTACGCAGCAGCC-3'<br>R-5'- GGTACCTACCCTCCCAGGGGTACAC-3' | 1350bp | (13) |

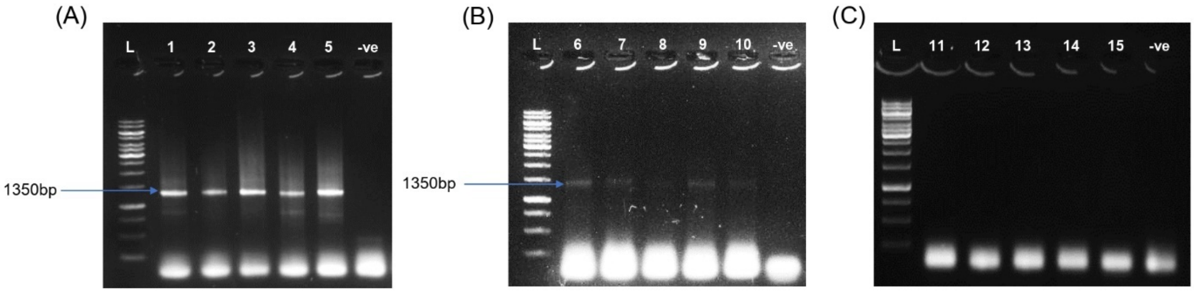


**Supplementary Figure S1. PCR Amplification of beta satellite DNA of CLCuV Infected Cotton Leaf Samples run on 1% Agarose gel.** (A) L = 1kb DNA ladder; Samples 1-5 (PFV-2 susceptible *G.hirsutum*); -ve (negative control with water) (B) L = 1kb DNA ladder; Samples 6-10 (PFV-1 partially tolerant *G.hirsutum*); -ve (negative control with water) (C) L = 1kb DNA ladder; Samples 11-15 (FDH-228 tolerant *G.arboreum*); -ve (negative control with water)

**Supplementary Table S2 -** Disease Severity Index Scale for Cotton Leaf Curl Disease (CLCuD) as per [14].

| **Disease Severity Index** | **Symptoms** |
|---|---|
| **0** | Complete absence of CLCuD symptoms and virus is undetectable in plant tissues using PCR. |
| **1** | Complete absence of symptoms, but virus can be detected in plant tissues using PCR. Thickening of veins or only presence of leaf enations on one or few leaves of upper canopy. |
| **2** | Thickening of small group of veins, no leaf curling, no reduction in leaf size and boll setting. No leaf curling observed. |
| **3** | Thickening of all veins, minor leaf curling and minor reduction in leaf size bu no reduction in boll setting. |
| **4** | Severe vein thickening in leaves, moderate leaf curling followed by minor deformity of internodes and minor reduction in leaf size and boll setting. |
| **5** | Severe vein thickening, moderate leaf curling and deformity of internodes with moderate reduction in leaf size and boll setting followed by moderate stunting of the cotton plant. |
| **6** | Severe vein thickening, leaf curling, reduction in leaf size, deformed internodes and stunting of the plant with no or few boll setting 6 >50 Highly susceptible |

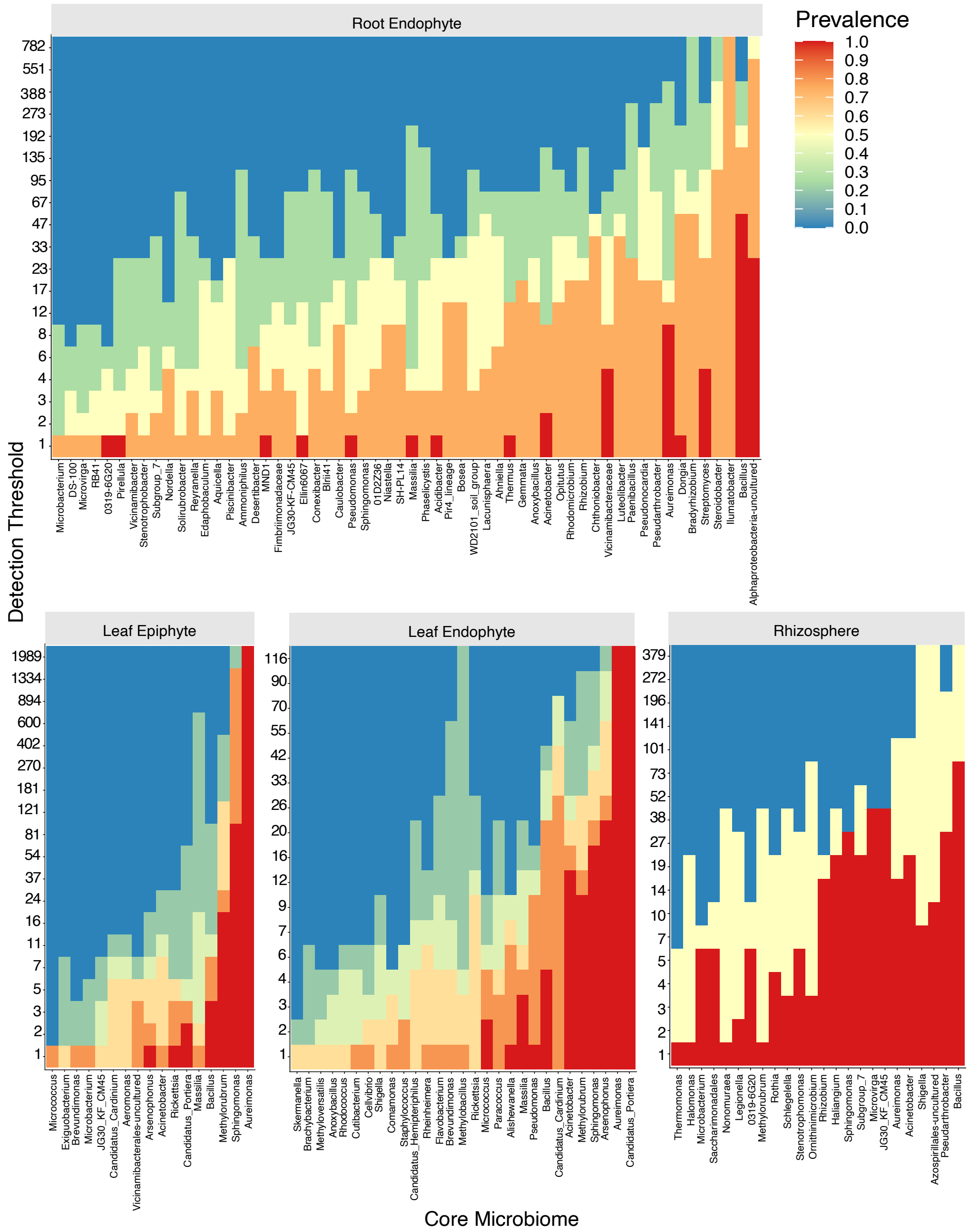


**Supplementary Figure S2. Core microbiome analysis *Gossypium hirsutum* PFV-2**: Heatmaps of the different meta-sample groups at genus level (Root Endophyte, Rhizosphere, Leaf Epiphyte, and Leaf Endophyte) derived from this study with a minimum prevalence of 50% and microbes sorted as left (low abundance) to right (high abundant) core microbiome.

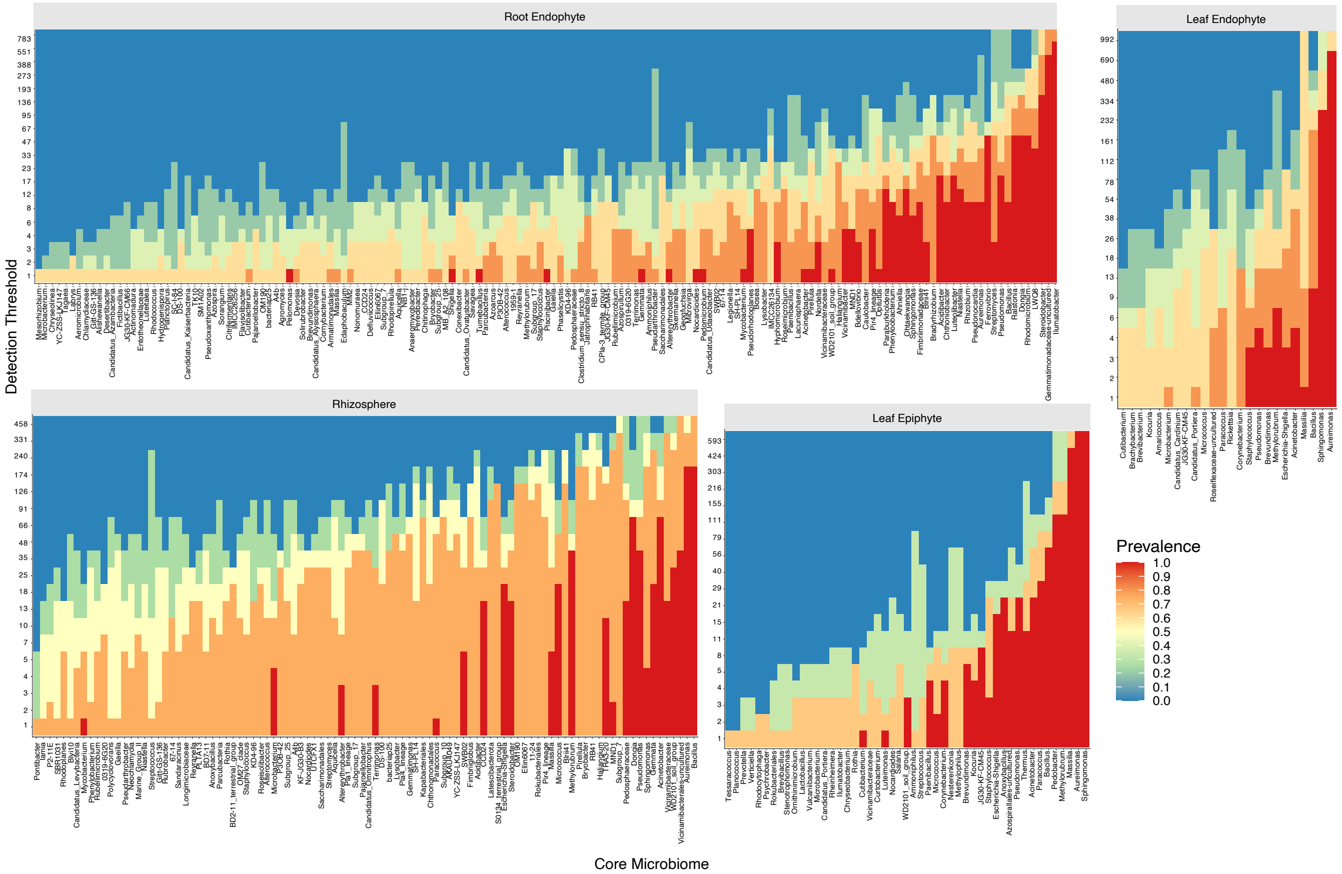


**Supplementary Figure S3. Core microbiome analysis of *Gossypium hirsutum* PFV1**: Heatmaps of the different meta-sample groups at genus level l (Root Endophyte, Rhizosphere, Leaf Epiphyte, and Leaf Endophyte) derived from this study with a minimum prevalence of 50% and microbes sorted as left (low abundance) to right (high abundant) core microbiome.

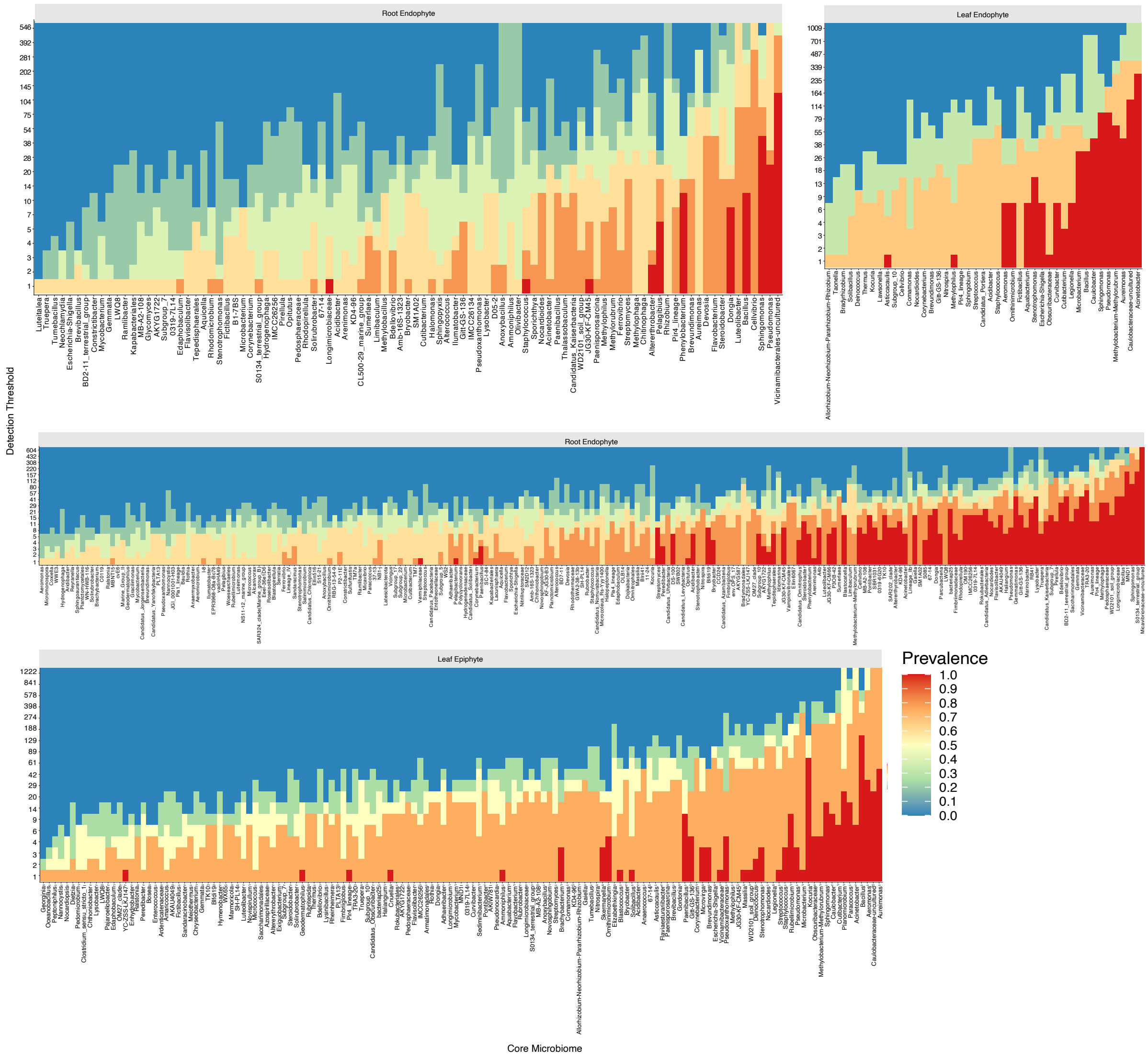


**Supplementary Figure S4. Core microbiome analysis of *Gossypium arboreum FDH228***: Heatmaps of the different meta-sample groups at genus level l (Root Endophyte, Rhizosphere, Leaf Epiphyte, and Leaf Endophyte) derived from this study with a minimum prevalence of 50% and microbes sorted as left (low abundance) to right (high abundant) core microbiome.

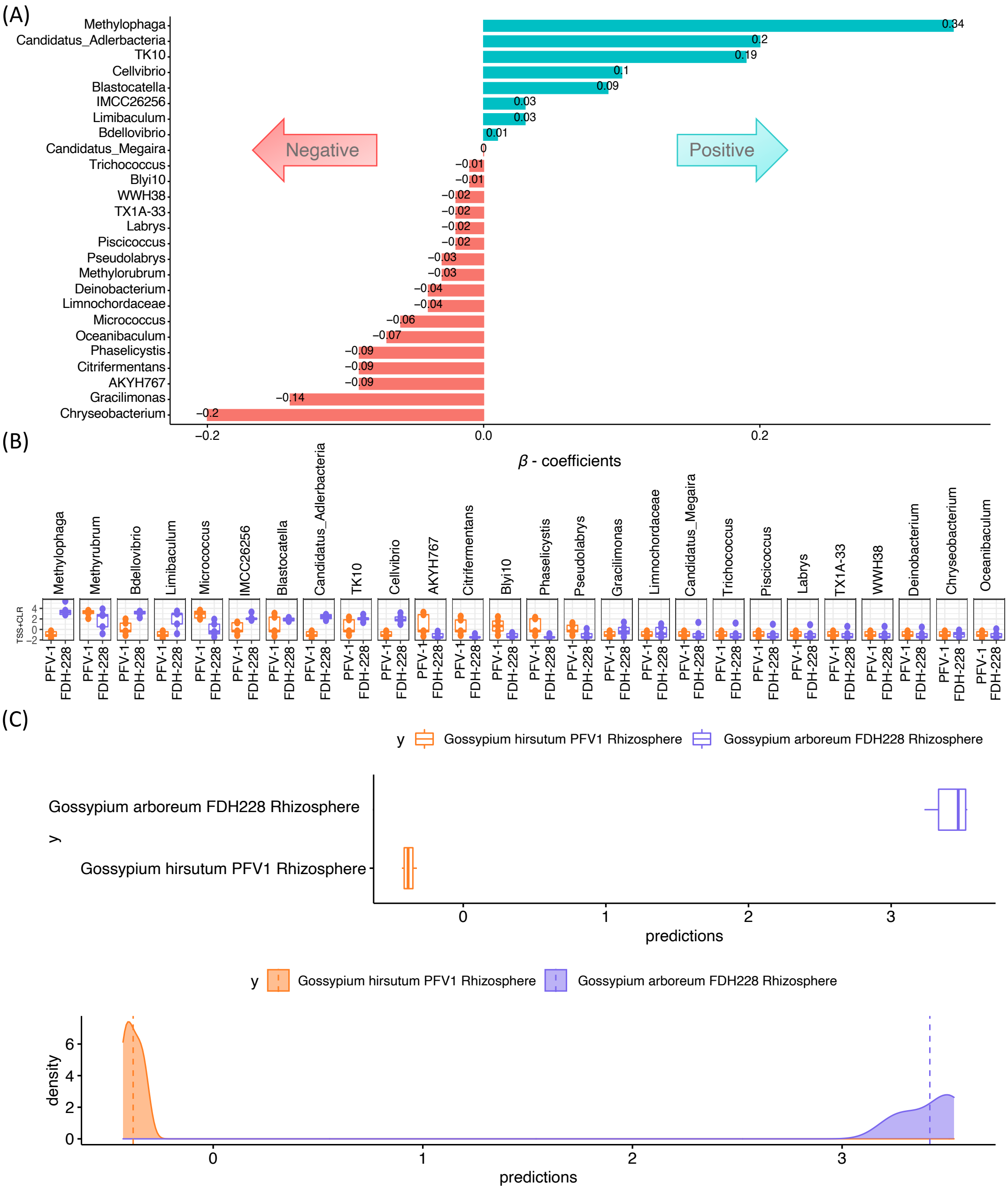


**Supplementary Figure S5. CODA LASSO regression for *Gossypium hirsutum* PFV1 and *Gossypium arboreum* FDH228 (Rhizosphere) for taxonomic abundance at Genus level**
A) $\beta$ −coefficients returned from CODA-LASSO procedure as two disjoint sets, with those that are associated with FDH228 (Positive), and those that are associated with PFV-1 (Negative); B) Expression levels of microbes selected from the procedure; and C) The density plot returned from the CODA-LASSO segregates the two groups provides a graphical assessment of the classification accuracy (top: true; bottom: predicted from the procedure).

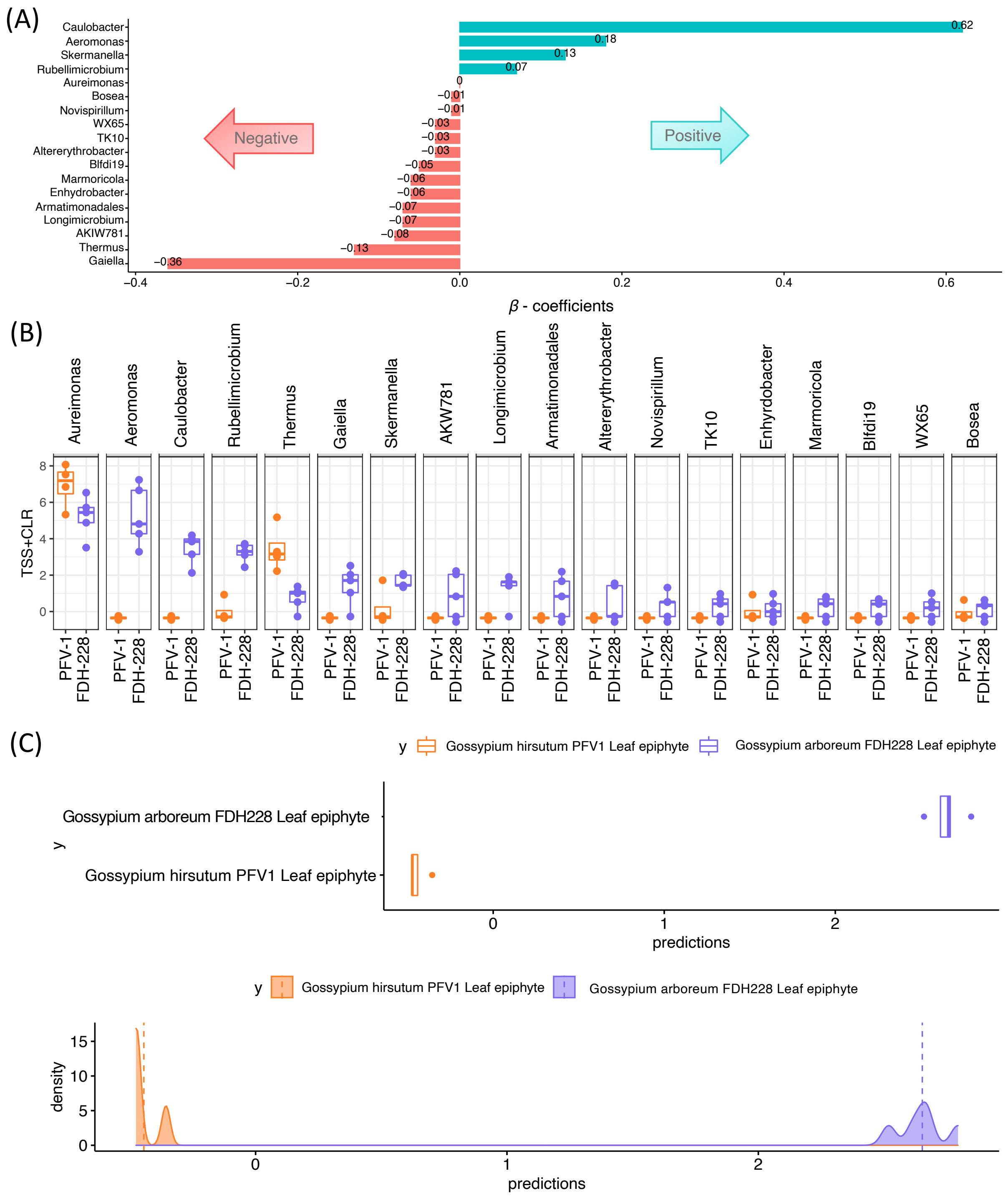


**Supplementary Figure S6. CODA LASSO regression for *Gossypium hirsutum* PFV1 and *Gossypium arboreum* FDH228 (Leaf epiphyte) for taxonomic abundance at Genus level**
A) $\beta$ −coefficients returned from CODA-LASSO procedure as two disjoint sets, with those that are associated with FH228 (Positive), and those that are associated with PFV-1 (Negative); B) Expression levels of microbes selected from the procedure; and C) The density plot returned from the CODA-LASSO segregates the two groups provides a graphical assessment of the classification accuracy (top: true; bottom: predicted from the procedure).

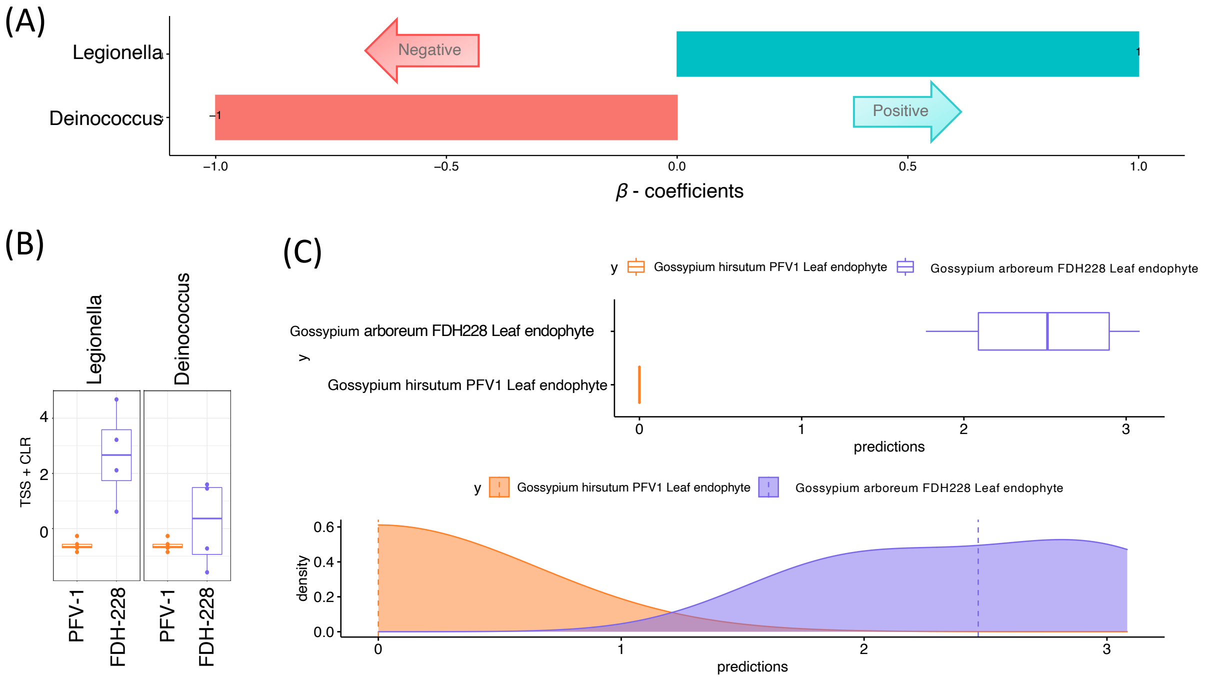


**Supplementary Figure S7. CODA LASSO regression for *Gossypium hirsutum* PFV1 and *Gossypium arboreum* FDH228 (Leaf endophyte) for taxonomic abundance at Genus level** A) $\beta$ −coefficients returned from CODA-LASSO procedure as two disjoint sets, with those that are associated with FDH228 (Positive), and those that are associated with PFV-1 (Negative); B) Expression levels of microbes selected from the procedure; and C) The density plot returned from the CODA-LASSO segregates the two groups provides a graphical assessment of the classification accuracy (top: true; bottom: predicted from the procedure).

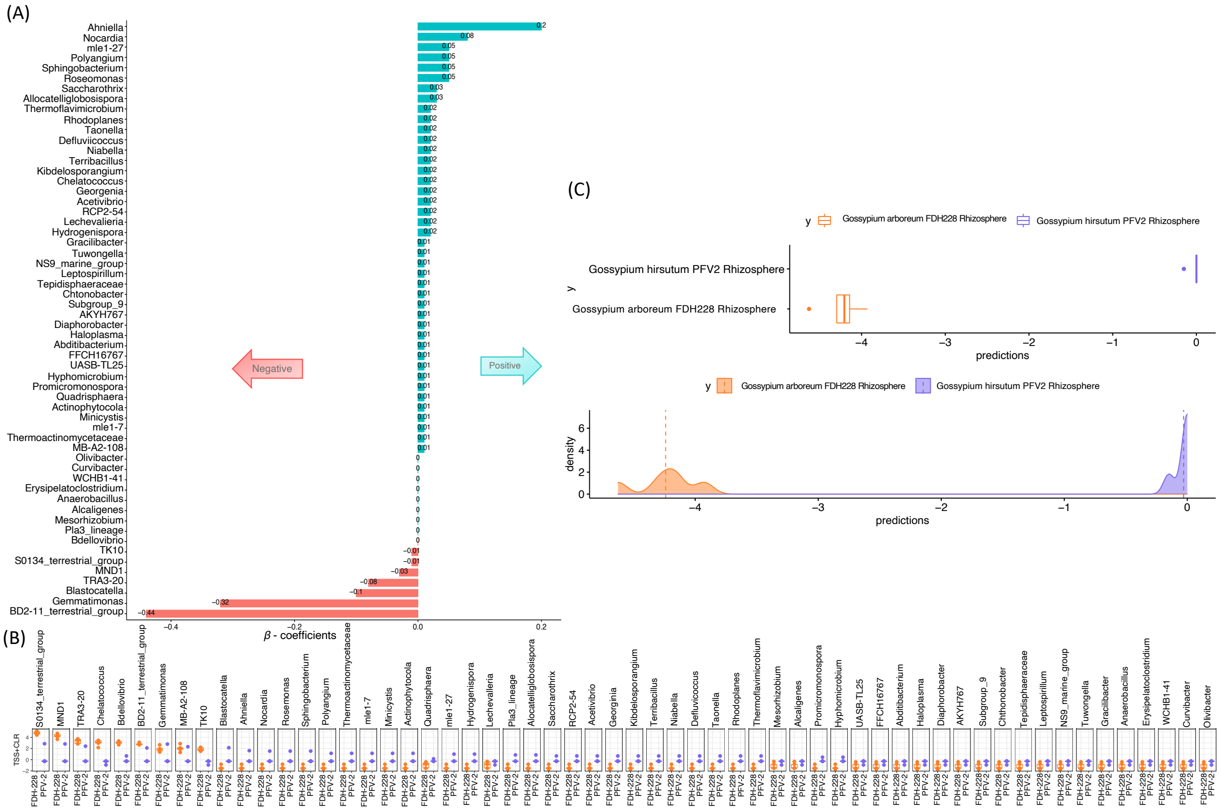

**Supplementary Figure S8. CODA LASSO regression for *Gossypium arboreum FDH-228* and *Gossypium hirsutum* PFV-2 (Rhizosphere) for taxonomic abundance at Genus level**
A) $\beta$ −coefficients returned from CODA-LASSO procedure as two disjoint sets (those that are positively related, and those that are negatively related with the temperature) B) The density plot returned from the CODA-LASSO segregates the two groups provides a graphical assessment of the classification accuracy (top: true; bottom: predicted from the procedure); C) Expression levels of microbes selected from the procedure.

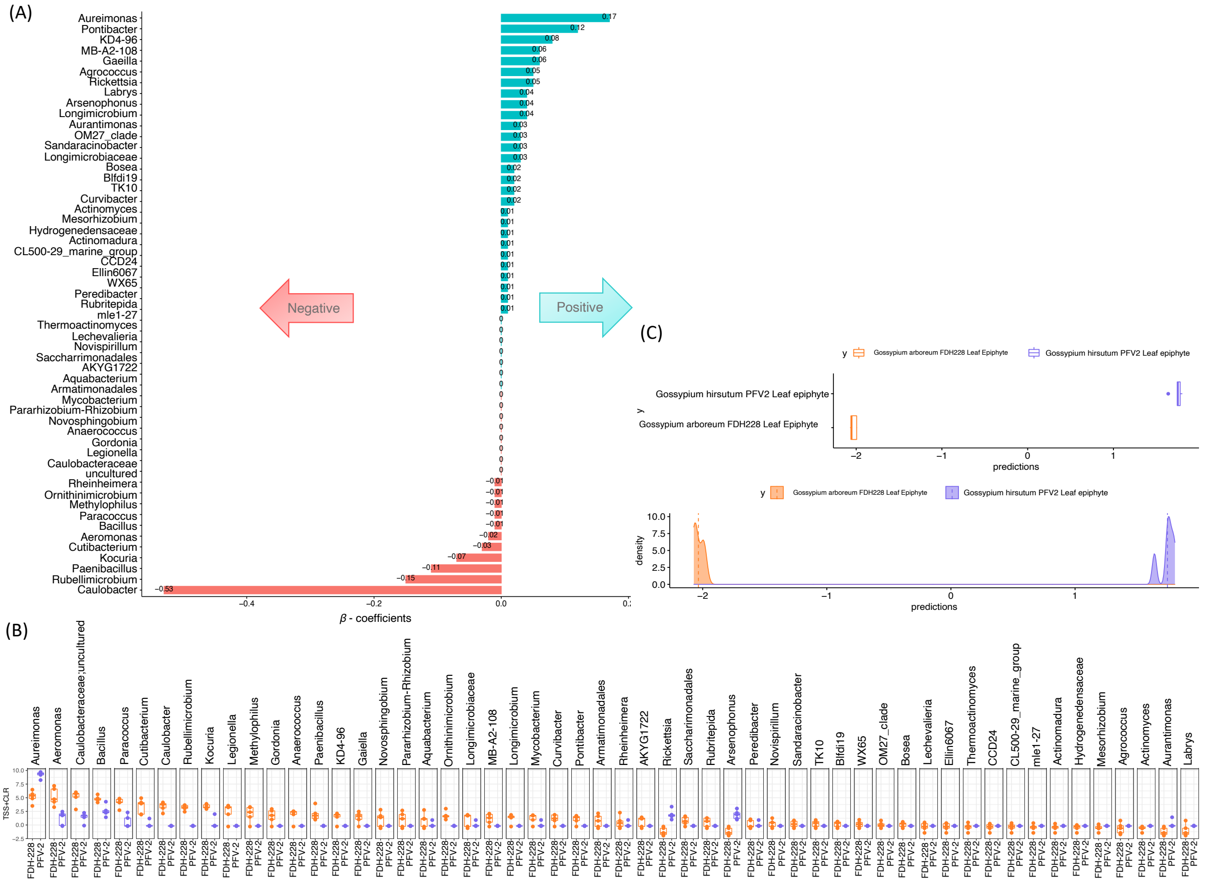


**Supplementary Figure S9. CODA LASSO regression for *Gossypium arboreum FDH-228* and *Gossypium hirsutum* PFV-2 (Leaf Epiphyte) for taxonomic abundance at Genus level** A) $\beta$ −coefficients returned from CODA-LASSO procedure as two disjoint sets (those that are positively related, and those that are negatively related with the temperature) B) The density plot returned from the CODA-LASSO segregates the two groups provides a graphical assessment of the classification accuracy (top: true; bottom: predicted from the procedure); C) Expression levels of microbes selected from the procedure.

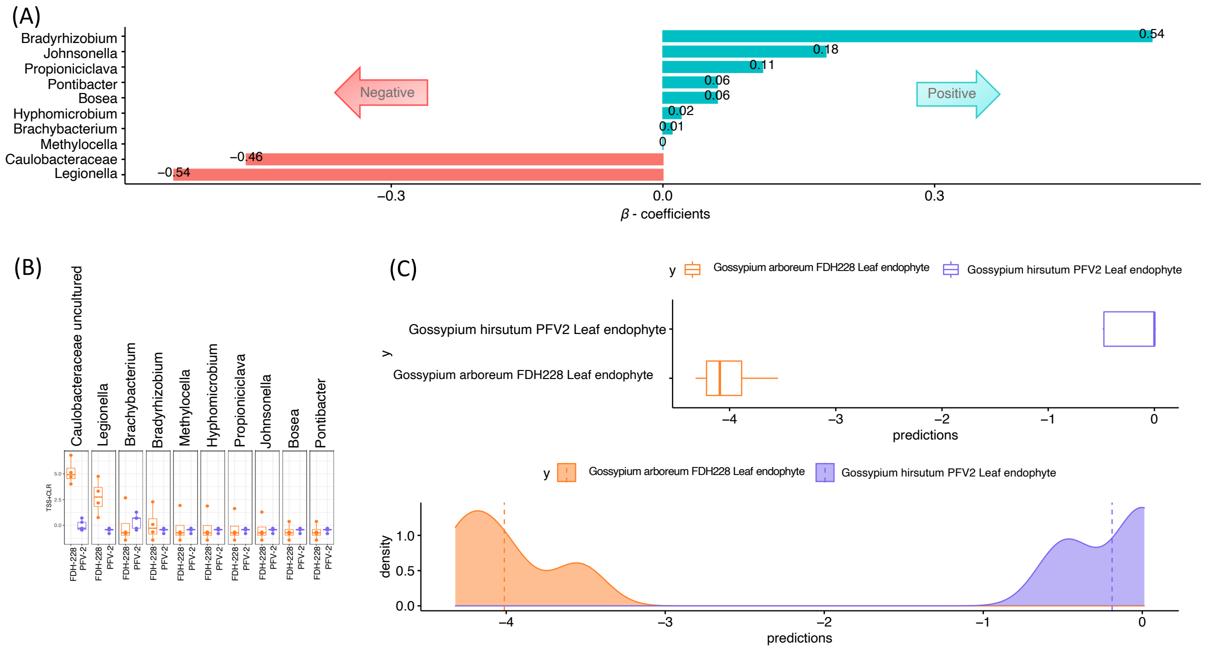


**Supplementary Figure S10. CODA LASSO regression for *Gossypium arboreum FDH-228* and *Gossypium hirsutum* PFV-2 (Leaf Endophyte) for taxonomic abundance at Genus level** A) $\beta$ −coefficients returned from CODA-LASSO procedure as two disjoint sets (those that are positively related, and those that are negatively related with the temperature) B) The density plot returned from the CODA-LASSO segregates the two groups provides a graphical assessment of the classification accuracy (top: true; bottom: predicted from the procedure); C) Expression levels of microbes selected from the procedure.

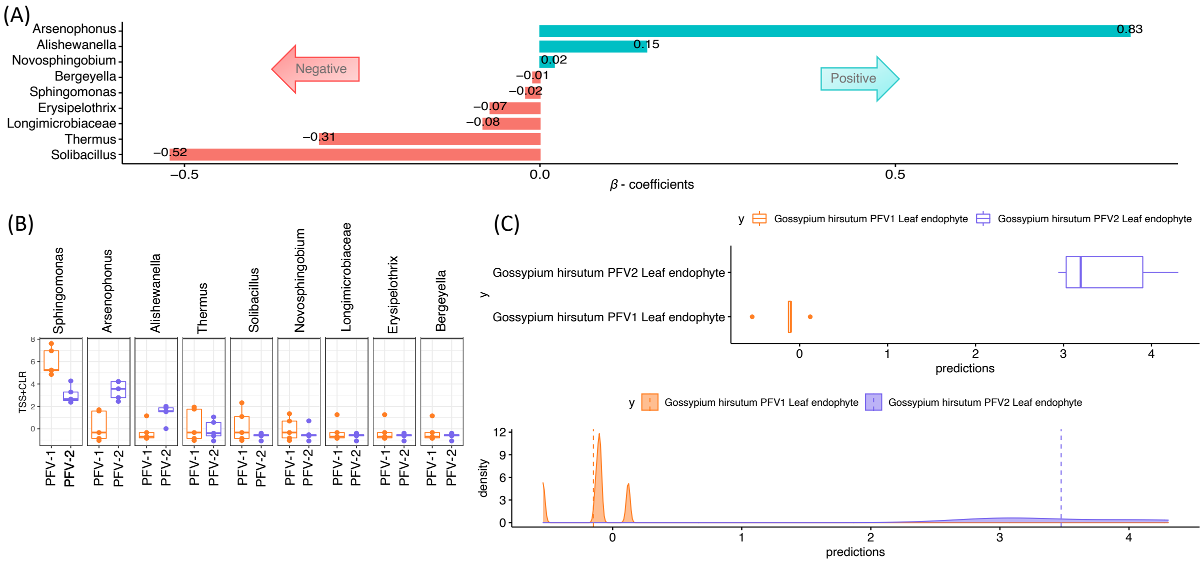


**Supplementary Figure S11. CODA LASSO regression for *Gossypium hirsutum* PFV-2 and *Gossypium hirsutum* PFV-1 (Leaf Endophyte)** A) $\beta$ −coefficients returned from CODA-LASSO procedure as two disjoint sets (those that are positively related, and those that are negatively related with the temperature) B) The density plot returned from the CODA-LASSO segregates the two groups provides a graphical assessment of the classification accuracy (top: true; bottom: predicted from the procedure); C) Expression levels of microbes selected from the procedure.

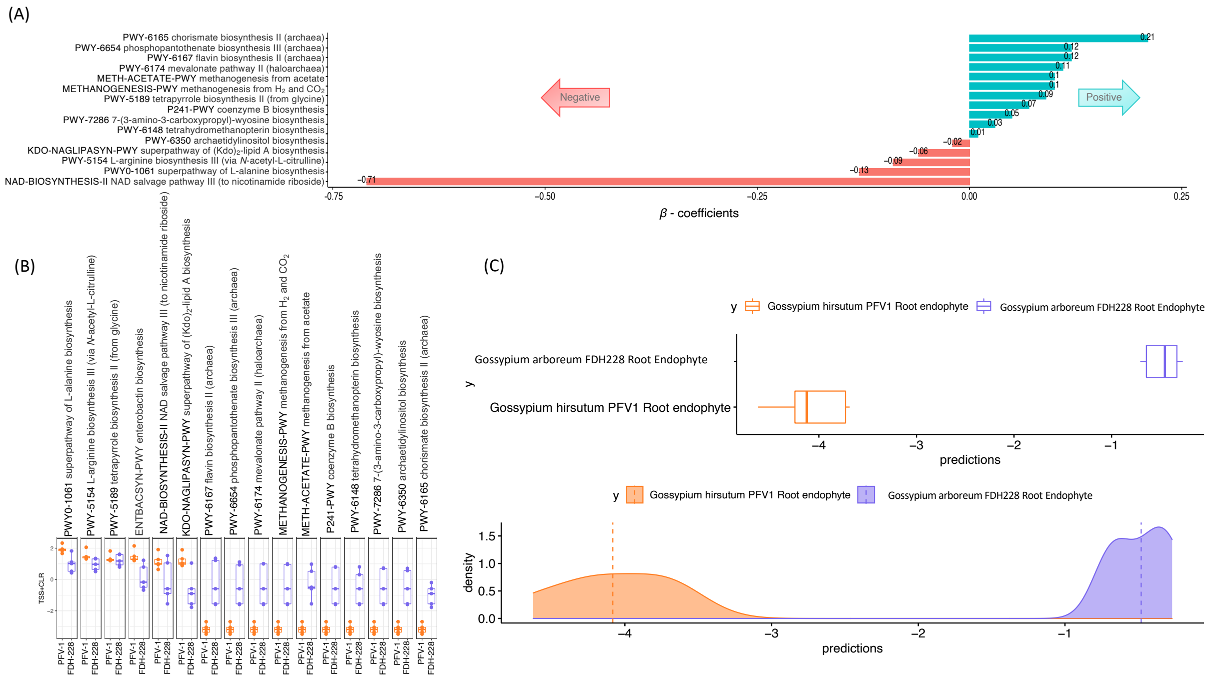


**Supplementary Figure S12. CODA LASSO regression for *Gossypium hirsutum* PFV-1 and *Gossypium arboreum FDH-228* (Root Endophyte) for MetaCyc pathways** A) $\beta$ −coefficients returned from CODA-LASSO procedure as two disjoint sets (those that are positively related, and those that are negatively related with the temperature) B) The density plot returned from the CODA-LASSO segregates the two groups provides a graphical assessment of the classification accuracy (top: true; bottom: predicted from the procedure); C) Expression levels of microbes selected from the procedure.

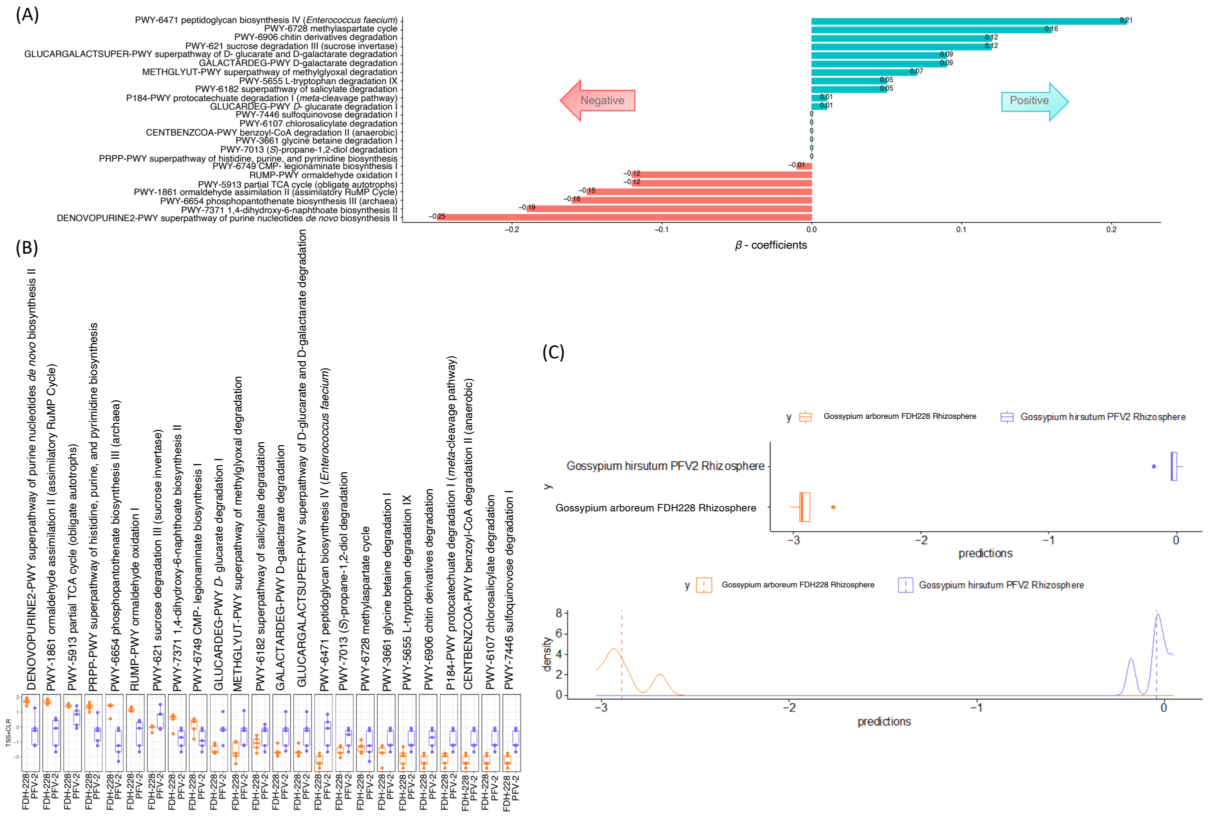

**Supplementary Figure S13. CODA LASSO regression for *Gossypium hirsutum* PFV-2 and *Gossypium arboreum FDH-228* (Rhizosphere) for MetaCyc pathways** A) $\beta$ −coefficients returned from CODA-LASSO procedure as two disjoint sets (those that are positively related, and those that are negatively related with the temperature) B) The density plot returned from the CODA-LASSO segregates the two groups provides a graphical assessment of the classification accuracy (top: true; bottom: predicted from the procedure); C) Expression levels of microbes selected from the procedure.

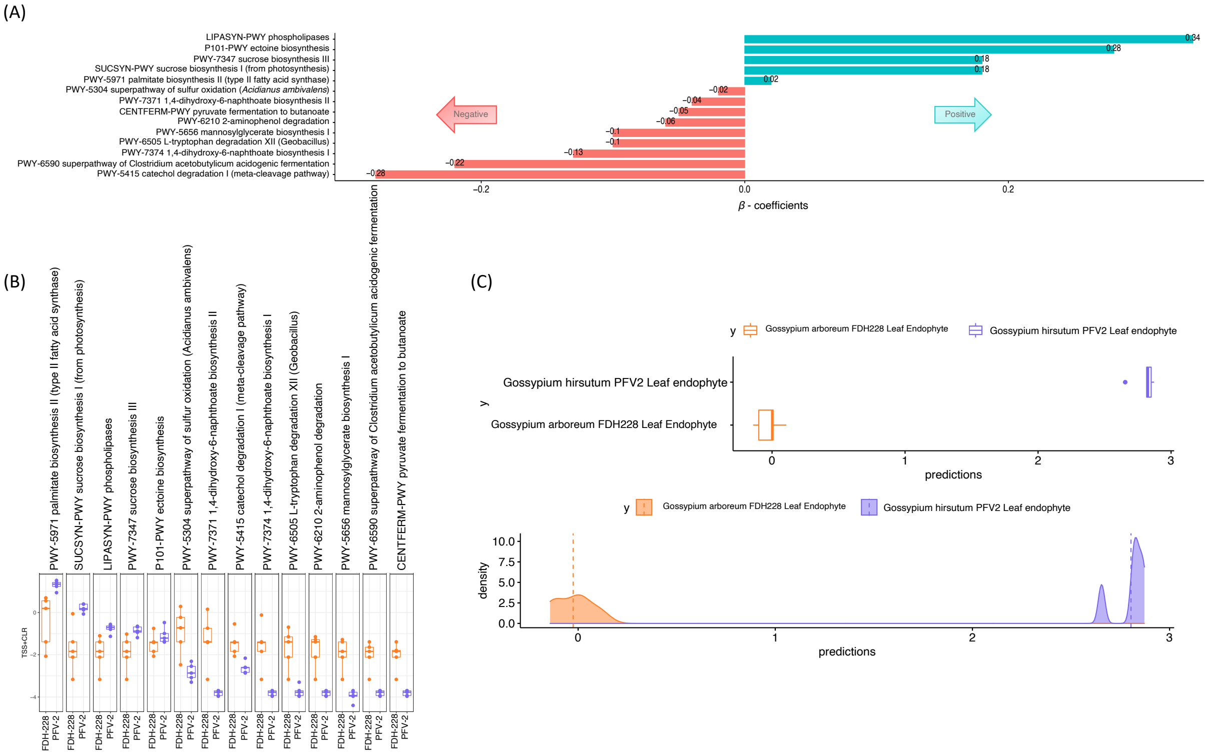


**Supplementary Figure S14. CODA LASSO regression for *Gossypium hirsutum* PFV-2 and *Gossypium arboreum FDH-228* (Leaf Endophyte) for MetaCyc pathways** A) $\beta$ −coefficients returned from CODA-LASSO procedure as two disjoint sets (those that are positively related, and those that are negatively related with the temperature) B) The density plot returned from the CODA-LASSO segregates the two groups provides a graphical assessment of the classification accuracy (top: true; bottom: predicted from the procedure); C) Expression levels of microbes selected from the procedure.

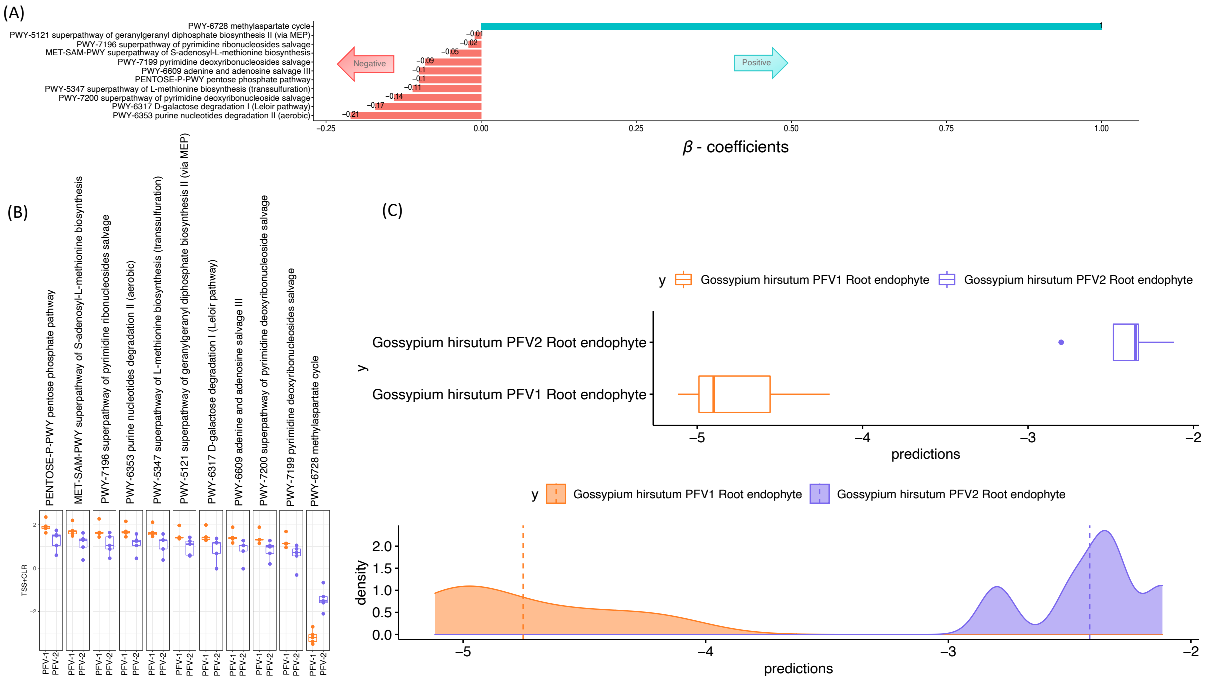


**Supplementary Figure S15. CODA LASSO regression for *Gossypium hirsutum* PFV-1 and *Gossypium hirsutum* PFV-2 (Root Endophyte) for MetaCyc pathways** A) $\beta$ –coefficients returned from CODA-LASSO procedure as two disjoint sets (those that are positively related, and those that are negatively related with the temperature) B) The density plot returned from the CODA-LASSO segregates the two groups provides a graphical assessment of the classification accuracy (top: true; bottom: predicted from the procedure); C) Expression levels of microbes selected from the procedure.

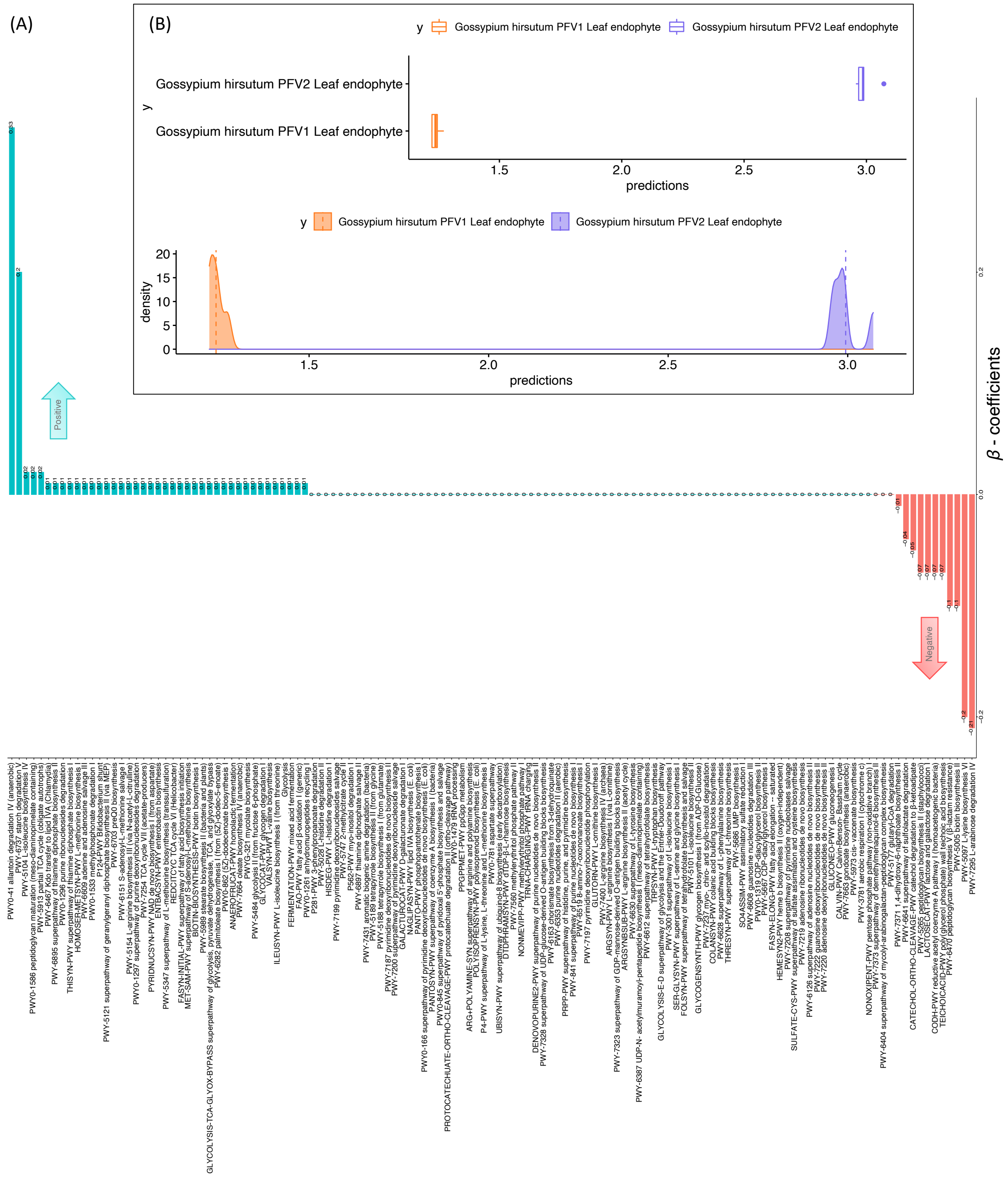


**Supplementary Figure S16. CODA LASSO regression for *Gossypium hirsutum* PFV-1 and *Gossypium hirsutum* PFV-2 (Leaf Endophyte) for MetaCyc pathways** A) $\beta$ −coefficients returned from CODA-LASSO procedure as two disjoint sets (those that are positively related, and those that are negatively related with the temperature) B) The density plot returned from the CODA-LASSO segregates the two groups provides a graphical assessment of the classification accuracy (top: true; bottom: predicted from the procedure).

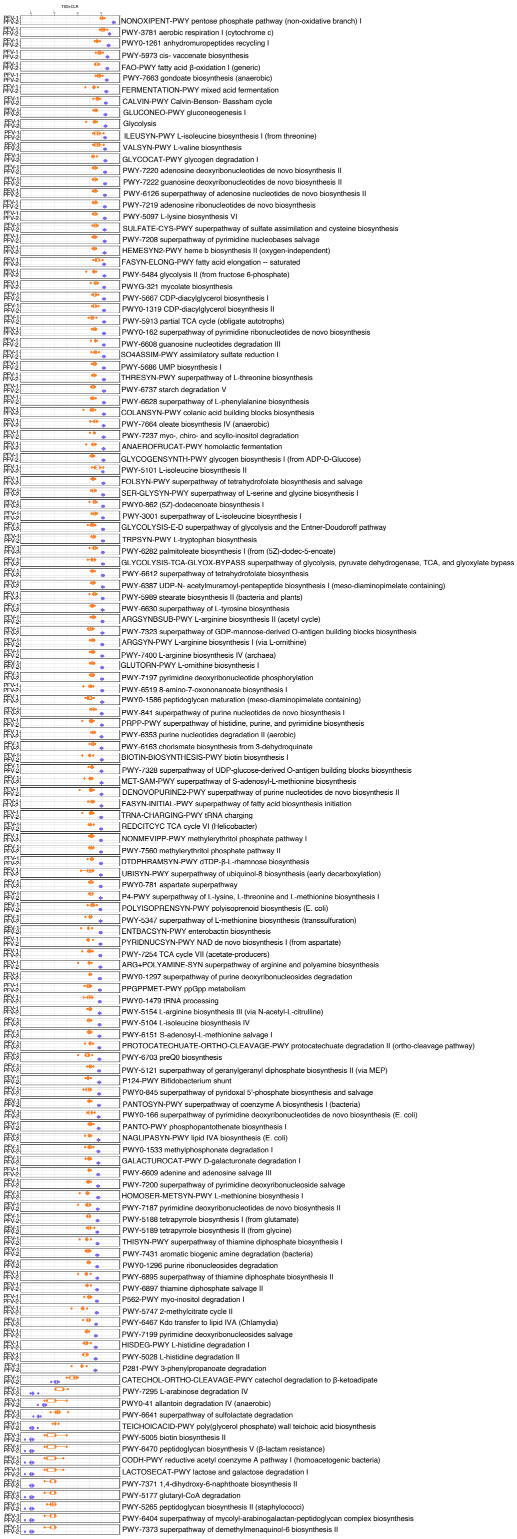


**Supplementary Figure S17. CODA LASSO regression for *Gossypium hirsutum* PFV-1 and *Gossypium hirsutum* PFV-2 (Leaf Endophyte) for MetaCyc pathways**. Expression levels of microbes selected from the procedure in Figure 2.5.5.

**Supplementary Table S3:** Top two most significantly positive and negative genera identified through CODA LASSO procedure in Supplementary Figures S5-S11 that compared compartments for different cotton varieties. Their significance in previously published literature is also shown. The variety in which the expression is higher is annotated with up arrow.

| Genus | Cotton Varieties Expression Trend | Compartment | Description |
|---|---|---|---|
| *Methylophaga* | PFV-1, FDH-228↑ | Rhizosphere | In root microbiome of healthy seagrass [15]<br>Isolated from rhizosphere of rice has PGPR activity [16] |
| *Candidatus adlerbacteria* | PFV-1, FDH-228 ↑ | Rhizosphere | *C. Adlerbacteria* in rhizosphere of Arecanut palms (Yellow Leaf Disease) [17] |
| *Gracilimonas* | PFV-1, FDH-228 ↑ | Rhizosphere | Dominant genus in halophyte [18]<br>Rhizosphere of halophytes [19] |
| *Chryseobacterium* | PFV-1, FDH-228 ↑ | Rhizosphere | Rhizosphere soil of Rhizoctonia solani Bare Patch Disease of Wheat [20] |
| *Caulobacter* | PFV-1, FDH-228 ↑ | Leaf Epiphyte | Root microbiomes of *Arabidopsis* [21], *Citrullus* [22],<br>Dioon [23] *Lavandula dentanta*<br>L. [24], *Populus deltoides* [25], and *Z. mays* [26, 27] |
| *Aeromonas* | PFV-1, FDH-228 ↑ | Leaf Epiphyte | Rhizosphere and endosphere of *Arabidopsis* [28] |
| *Thermus* | PFV-1, FDH-228 ↓ | Leaf Epiphyte | Apple flower microbiome [29] |
| *Gaiella* | PFV-1, FDH-228 ↑ | Leaf Epiphyte | Rhizosphere of resistant strawberry cultivars against soil-borne fungal pathogens [30] |
| *Legionella* | PFV-1, FDH-228 ↑ | Leaf endophyte | Leaf microbiome of radish (*Raphanus sativus*), lettuce (*Lactuca sativa*), and pakchoi (*Brassica chinensis*) [31] |

| | | | |
|---|---|---|---|
| *Deinococcus* | PFV-1, FDH-228 ↑ | Leaf endophyte | Native-grown *N. attenuata* plant roots [32]<br>Tomato leaf microbiome [33]<br>Pear and apple bark [34] |
| *Ahniella* | FDH-228, PFV-2 ↑ | Rhizosphere | Isolated from sandy soil near a stream [35] |
| *Nocardia* | FDH-228, PFV-2 ↑ | Rhizosphere | Isolated from a root nodule of an Alnus glutinosa plant growing in Leazes Park, Newcastle upon Tyne, UK [36]<br>Rhizosphere of desert plant *Calotropis procera* [37] |
| *Gemmatimonas* | PFV-2, FDH-228 ↑ | Rhizosphere | Rhizosphere of mono-cropped system [38]<br>Rhizosphere microbiome of Jerusalem artichoke [39]<br>Anaerobic–aerobic sequential batch reactor [40]<br>Rhizosphere microbiome of Rhizoma Alismatis (Chinese medicinal herb) [41]<br>Rotation soil of chilli pepper-banana [42]<br>Surface water of a stream in the Zackenberg Valley in High Arctic Greenland [43]<br>Freshwater Swan Lake in the western Gobi Desert [44] |
| *BD2-11_terrestrial group* | PFV-2, FDH-228 ↑ | Rhizosphere | Hypersaline sodalakes [45] |
| *Aureimonas* | FDH-228, PFV-2 ↑ | Leaf Epiphyte | Endorhizosphere of sunflower plants [46]<br>Rice seed microbiome [47, 48] |

| | | | |
|---|---|---|---|
| *Pontibacter* | PFV-2, FDH-228 ↑ | Leaf Epiphyte | Saline-sandy area soil microbiome [49] |
| *Rubellimicrobium* | PFV-2, FDH-228 ↑ | Leaf Epiphyte | Saline-sandy area soil microbiome [49] |
| *Bradyrhizobium* | PFV-2, FDH-228 ↑ | Leaf endophyte | Soybean rhizosphere [50]<br>Canola rhizosphere [51]<br>Metarhizium treated root tissue of beans [52] |
| *Johnsonella* | FDH-228, PFV-2 ↑ | Leaf endophyte | Not available for plant microbiome |
| *Caulobacteraceae* | PFV-2, FDH-228 ↑ | Leaf endophyte | Rhizosphere microbiome of Solanaceae eggplant resistant varieties against bacterial wilt resistance [53] |
| *Arsenophonus* | PFV-1, PFV-2 ↑ | Leaf endophyte | Pesticide treated tea leaf microbiome [54]<br>Insect pathogen [55] |
| *Alishewanella* | PFV-1, PFV-2 ↑ | Leaf endophyte | Core pine nut microbiome [56] |
| *Solibacillus* | PFV-2, PFV-1 ↑ | Leaf endophyte | Cowpea soil microbiome [57]<br>Healthy tomato rhizosphere soil [58] |
| *Longimicrobiaceae* | PFV-2, PFV-1 ↑ | Leaf endophyte | Potato rhizosphere microbiome [59, 60] |
| *Bdellovibrio* | PFV-2, FDH-228 ↑ | Rhizosphere | Citrus root-associated microbiome [61]<br>Rhizosphere of resistant variety [62] |

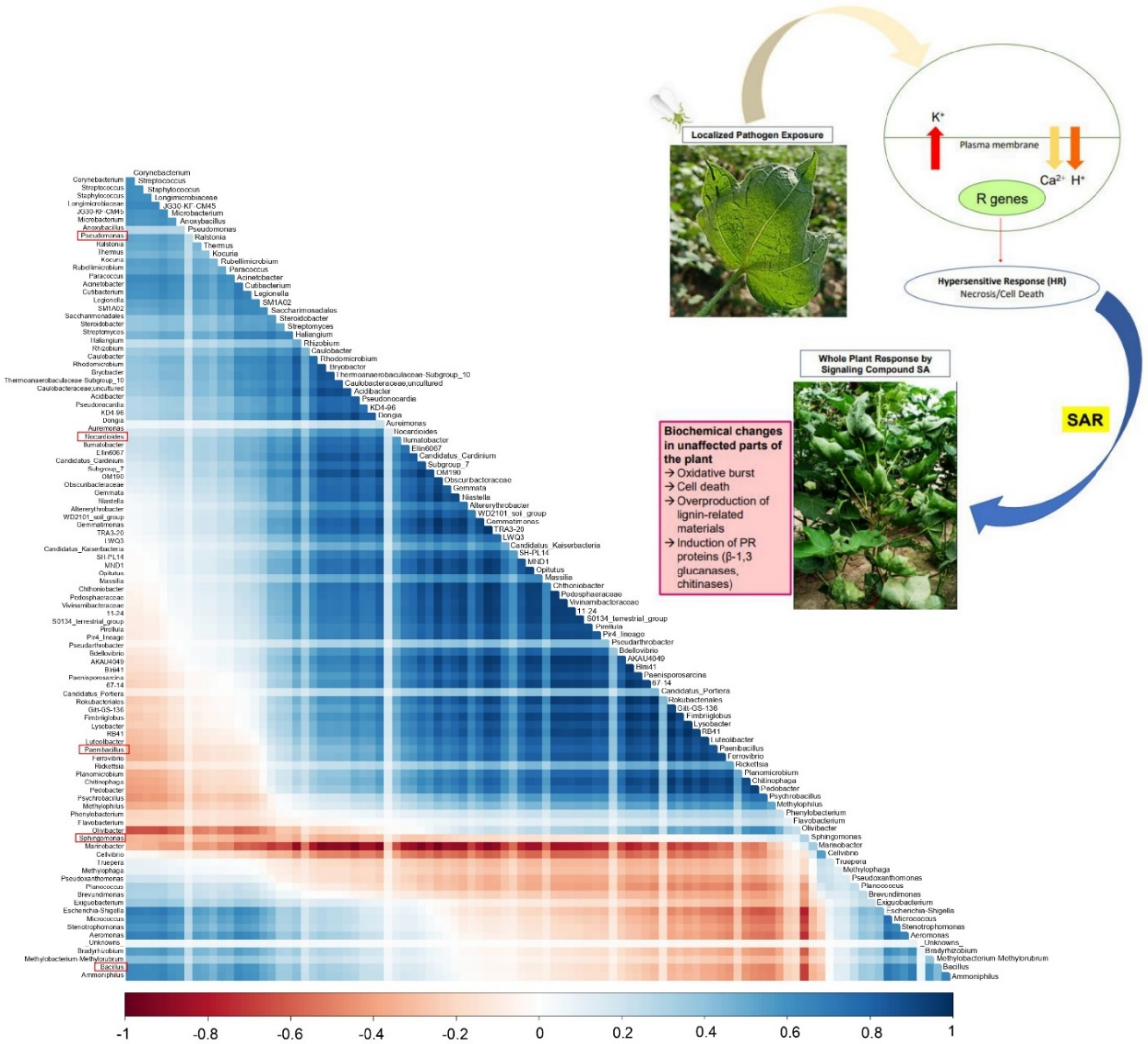


**Supplementary Figure S18. Co-occurrence relationship between microbes recovered from the residual covariance matrix $\Sigma$ that are not explained by the environmental covariates in the GLLVM model in Figure 4.** Here, blue represent the positive correlation, and red represent the negative relationship. The highlighted bacteria are capable of SA production.

**Supplementary Table S4: Bacterial Strains isolated from different plant compartments of CLCuD susceptible, partially tolerant, and tolerant cotton varieties. This** table shows their isolate codes, morphological characteristics, indole acetic acid and salicylic acid production results.

| Plant Variety | Plant Compartment | Isolate Code | Shape | Colour | Elevation | Margin | Appearance | SA production | IAA production | 16S rRNA based identification |
|---|---|---|---|---|---|---|---|---|---|---|
| PFV-2 | Leaf epiphyte | V2 lep 10-8 | Circular | white | flat | entire | opaque glossy | X | ✓ | - |
| | | V2 lep 10-10 | circular | dark pink | flat | entire | very small | X | X | - |
| | | V2 lep 10-6 | Circular | yellow | raised | entire | smooth, transparent | X | ✓ | - |
| | Leaf endophyte | V2 len 3-2 | Circular | white | raised | entire | small, opaque | X | ✓ | - |
| | | V2 len 3-3 | Circular | neon lime | flat | entire | small, translucent | X | ✓ | - |
| | | V2 len 10-6 | Circular | yellow | flat | entire | transparent | X | ✓ | - |
| | | V2 len 10-5 | Circular | yellow | raised | entire | transparent | X | ✓ | - |
| | | V2 len 10-2 | Irregular | Dark yellow | flat | undulate | opaque | X | ✓ | - |
| | | V2 len 6-1 | Circular | white | flat | entire | small | X | ✓ | - |
| | Rhizosphere | V2 rhi 1-1 | Circular | White | raised | Entire | Opaque glossy | X | ✓ | - |
| | | V2 rhi 7-1 | Irregular | Off white | flat | Undulate | opaque | X | ✓ | - |
| | Root endophyte | V2 ren 3-2 | Filiform | Translucent white | flat | undulate | very small | X | X | - |
| PFV-1 | Leaf epiphyte | EPI.PFV1.P1.TSA.C1 | Circular | Yellow Inner | raised | entire | Small | X | ✓ | - |
| | | EPI.PFV1.P3.TSA.C2 | Irregular | Light yellow | convex | umbonate | Small | X | ✓ | - |
| | | EPI.PFV1.P3.TSA.C1 | Circular | Orange | convex | entire | Large | X | ✓ | - |

| | | | | | | | | | | |
|---|---|---|---|---|---|---|---|---|---|---|
| | Leaf endophyte | ENDO.PFV1.P1.TSA.C1 | Irregular | Pure white | flat | umbonate | Large | X | √ | - |
| | | ENDO.PFV1.P3.TSA.C3 | Circular | Off white outer | raised | entire | Small | X | √ | - |
| | | ENDO.PFV1.P1.TSA.C2 | Circular | Dull white | raised | entire | Small | X | √ | - |
| | | ENDO.PFV1.P3.TSA.C2 | Circular | Cream white | convex | entire | small | X | √ | - |
| | | Rh.pfv1.P9.TSA.C2 | Circular | White | Convex | Entire | Smooth | X | √ | - |
| | | Rh.PFV1.P6.TSA.C1 | Circular | Yellow | Flat | Entire | Viscid | X | √ | - |
| | | RT En.PFV1.P3.TSA.C3 | Irregular | Milky white | Convex | Undulate | Viscid | X | √ | - |
| | | RT En PFV1.P3.TSA.C1 | Irregular | Off white | Flat | Undulate | Smooth | X | √ | - |
| FDH-228 | Leaf epiphyte | FHAP2.6 | Circular | Pink | Convex | Entire | smooth,moist & coccibacilus | √ | √ | uncultured *Serratia spp.* |
| | | FHAP7.7 | Circular | Yellow | Convex | Entire | smooth,moist & rods | √ | √ | *Bacillus spp.* |
| | | FHAP1.8 | Circular | Off white | Raised | Entire | glistery,dry & rods | X | √ | - |
| | Leaf endophyte | FHAN2.7 | Circular | Lemon | Convex | Entire | Glistery, moist & rod | X | √ | - |
| | | FHAN2.8 | Circular | Milky | Convex | Entire | Smooth, dry & rod | √ | X | *Fictibacillus spp.* |
| | Rhizosphere | Rh. P1. -5. C1 | Circular | Yellow | Raised | Entire | Small | X | X | - |
| | | Rh. P1. -5. C2 | Circular | Pure White | Raised | Entire | Small | X | X | - |

**Supplementary Table S5:** Soil Classification and Chemical Characteristics

| Soil Characteristic | Values |
|---|---|
| **pH** | **7.43** |
| **EC (µScm$^{-1}$)** | **99.52** |
| **Organic matter (%)** | **3.140** |
| **Total Kjeldhal Nitrogen (%)** | **0.028** |
| **Total Phosphorus $P_2O_5$ (%)** | **0.070** |
| **Total Potassium (%)** | **0.290** |
| **Total Calcium (%)** | **0.450** |
| **Total Magnesium (%)** | **0.380** |
| **Total Sodium (%)** | **8.710** |
| **Total Manganese (mg/Kg)** | **12.50** |
| **Soil texture** | **Sandy Loam** |

References:


[1] Amir, A., McDonald, D., Navas-Molina, J. A., Kopylova, E., Morton, J. T., Zech Xu, Z., ... & Knight, R. (2017). Deblur rapidly resolves single-nucleotide community sequence patterns. MSystems, 2(2), e00191-16.]

[2] Douglas, G. M., Maffei, V. J., Zaneveld, J. R., Yurgel, S. N., Brown, J. R., Taylor, C. M., ... & Langille, M. G. (2020). PICRUSt2 for prediction of metagenome functions. Nature biotechnology, 38(6), 685-688.]

[3] Oksanen, J., Blanchet, F. G., Kindt, R., Legendre, P., Minchin, P. R., O'hara, R. B., ... & Oksanen, M. J. (2013). Package 'vegan'. Community ecology package, version, 2(9), 1-295.

[4] McMurdie, P. J., & Holmes, S. (2013). phyloseq: an R package for reproducible interactive analysis and graphics of microbiome census data. PloS one, 8(4), e61217.

[5] Zhang, Y., Jing, G., Chen, Y., Li, J., & Su, X. (2021). Hierarchical Meta-Storms enables comprehensive and rapid comparison of microbiome functional profiles on a large scale using hierarchical dissimilarity metrics and parallel computing. Bioinformatics Advances, 1(1), vbab003.

[6] Lahti, L., Shetty, S., Blake, T., & Salojarvi, J. (2017). Tools for microbiome analysis in R. Version, 1(5), 28. https://bioconductor.org/packages/release/bioc/html/microbiome.html

[7] Susin, A., Wang, Y., Lê Cao, K. A., & Calle, M. L. (2020). Variable selection in microbiome compositional data analysis. NAR Genomics and Bioinformatics, 2(2), lqaa029

[8] Lu, J., Shi, P., & Li, H. (2019). Generalized linear models with linear constraints for microbiome compositional data. Biometrics, 75(1), 235-244.

[9] Calle, M., & Susin, T. (2022). coda4microbiome: Compositional Data Analysis for Microbiome Studies. R package version 0.1.1, https://CRAN.R-project.org/package=coda4microbiome.

[10] Niku, J., Hui, F. K., Taskinen, S., & Warton, D. I. (2019). gllvm: Fast analysis of multivariate abundance data with generalized linear latent variable models in r. Methods in Ecology and Evolution, 10(12), 2173-2182.

[11] Colco, R. (2005). "Gram Staining". Current Protocols in Microbiology. Appendix 3 (1): Appendix 3C.

[12] Herlemann, D. P., Labrenz, M., Jürgens, K., Bertilsson, S., Waniek, J. J., & Andersson, A. F. (2011). Transitions in bacterial communities along the 2000 km salinity gradient of the Baltic Sea. The ISME journal, 5(10), 1571-1579.

[13] Monga, D., Kumar, R., & Kumar, M. (2005). Detection of DNA-A and satellite (DNA-β) in cotton leaf curl virus (CLCuV) infected weeds and cotton plants using PCR technique. Journal of Cotton Research and Development, *19*(1), 105-108.

[14] Akhtar, K. P., Haidar, S., Khan, M. K. R., Ahmad, M., Sarwar, N., Murtaza, M. A., & Aslam, M. (2010). Evaluation of *Gossypium* species for resistance to cotton leaf curl Burewala virus. Annals of applied biology, 157(1), 135-147.

[15] Martin, B. C., Alarcon, M. S., Gleeson, D., Middleton, J. A., Fraser, M. W., Ryan, M. H., ... & Kilminster, K. (2020). Root microbiomes as indicators of seagrass health. FEMS Microbiology Ecology, 96(2), fiz201.

[16] Bal, H. B., & Adhya, T. K. (2021). Alleviation of submergence stress in rice seedlings by plant growth-promoting rhizobacteria with ACC deaminase activity. Frontiers in Sustainable Food Systems, 5, 606158.

[17] Paulraj, S., Bhat, R., Rajesh, M. K., Ramesh, S.V., Priya, U.K., Pandian, T. P. R., Hegde, V., and Chowdappa, P. 2021. Microbiome-mediated Rhizosphere Nitrogen Transformation Cycle (RNTC) potentially underlies the disease severity in Arecanut Yellow Leaf Disease (YLD): Insights from metagenomics. In: PLACROSYM XXIV (Eds.) Dhanapal, K., Kumar, K. P., Shadanaika, Ali, M.A.A., Varghese, J. J., Saju, K.A., Oommen, M., and Thiyagarajan, P., Indian Cardamom Research Institute/Indian Society for Plantation Crops, Kasaragod, Kerala. pp. 221-222.

[18] Gao, L., Huang, Y., Liu, Y., Mohamed, O. A. A., Fan, X., Wang, L., ... & Ma, J. (2022). Bacterial community structure and potential microbial coexistence mechanism associated with three halophytes adapting to the extremely hypersaline environment. Microorganisms, 10(6), 1124.

[19] Li, Y., Kong, Y., Teng, D., Zhang, X., He, X., Zhang, Y., & Lv, G. (2018). Rhizobacterial communities of five co-occurring desert halophytes. PeerJ, 6, e5508.

[20] Yin, C., Hulbert, S. H., Schroeder, K. L., Mavrodi, O., Mavrodi, D., Dhingra, A., ... & Paulitz, T. C. (2013). Role of bacterial communities in the natural suppression of Rhizoctonia solani bare patch disease of wheat (Triticum aestivum L.). Applied and Environmental Microbiology, 79(23), 7428-7438.

[21] Lundberg, D. S., Lebeis, S. L., Paredes, S. H., Yourstone, S., Gehring, J., Malfatti, S., ... & Dangl, J. L. (2012). Defining the core Arabidopsis thaliana root microbiome. Nature, 488(7409), 86-90.

[22] Ren, R., Yang, X., Xu, J., Zhang, M., Liu, G., & Yao, X. (2019). Genome-wide identification and analysis of GDSL-type esterases/lipases in watermelon (Citrullus lanatus).

[23] Gutiérrez-García, K., Bustos-Díaz, E. D., Corona-Gómez, J. A., Ramos-Aboites, H. E., Sélem-Mojica, N., Cruz-Morales, P., ... & Cibrián-Jaramillo, A. (2019). Cycad coralloid roots contain bacterial communities including cyanobacteria and Caulobacter spp. that encode niche-specific biosynthetic gene clusters. Genome Biology and Evolution, 11(1), 319-334.

[24] Pereira, S. I. A., Monteiro, C., Vega, A. L., & Castro, P. M. (2016). Endophytic culturable bacteria colonizing Lavandula dentata L. plants: isolation, characterization and evaluation of their plant growth-promoting activities. Ecological Engineering, 87, 91-97.

[25] Brown, S. D., Klingeman, D. M., Lu, T. Y. S., Johnson, C. M., Utturkar, S. M., Land, M. L., ... & Pelletier, D. A. (2012). Draft genome sequence of Rhizobium sp. strain PDO1-076, a bacterium isolated from Populus deltoides.

[26] Naveed, M., Mitter, B., Reichenauer, T. G., Wieczorek, K., & Sessitsch, A. (2014). Increased drought stress resilience of maize through endophytic colonization by Burkholderia phytofirmans PsJN and Enterobacter sp. FD17. Environmental and Experimental Botany, 97, 30-39.

[27] Gao, J., Luo, M., Peng, H., Chen, F., & Li, W. (2019). Characterization of cadmium-responsive MicroRNAs and their target genes in maize (Zea mays) roots. BMC Molecular Biology, 20(1), 1-9.

[28] He, D., Singh, S. K., Peng, L., Kaushal, R., Vílchez, J. I., Shao, C., ... & Zhang, H. (2022). Flavonoid-attracted Aeromonas sp. from the Arabidopsis root microbiome enhances plant dehydration resistance. The ISME Journal, 16(11), 2622-2632.

[29] Shade, A., McManus, P. S., & Handelsman, J. (2013). Unexpected diversity during community succession in the apple flower microbiome. MBio, 4(2), e00602-12.

[30] Lazcano, C., Boyd, E., Holmes, G., Hewavitharana, S., Pasulka, A., & Ivors, K. (2021). The rhizosphere microbiome plays a role in the resistance to soil-borne pathogens and nutrient uptake of strawberry cultivars under field conditions. Scientific Reports, 11(1), 1-17.

[31] Li, W. J., Li, H. Z., An, X. L., Lin, C. S., Li, L. J., & Zhu, Y. G. (2022). Effects of manure fertilization on human pathogens in endosphere of three vegetable plants. Environmental Pollution, 314, 120344.

[32] Santhanam, R., Oh, Y., Kumar, R., Weinhold, A., Luu, V. T., Groten, K., & Baldwin, I. T. (2017). Specificity of root microbiomes in native‐grown Nicotiana attenuata

and plant responses to UVB increase Deinococcus colonization. Molecular Ecology, 26(9), 2543-2562.

[33] Toju, H., Okayasu, K., & Notaguchi, M. (2019). Leaf-associated microbiomes of grafted tomato plants. Scientific reports, 9(1), 1-11.

[34] Arrigoni, E., Antonielli, L., Pindo, M., Pertot, I., & Perazzolli, M. (2018). Tissue age and plant genotype affect the microbiota of apple and pear bark. Microbiological research, 211, 57-68.

[35] Hwang, W. M., Ko, Y., Kim, J. H., & Kang, K. (2018). Ahniella affigens gen. nov., sp. nov., a gammaproteobacterium isolated from sandy soil near a stream. International Journal of Systematic and Evolutionary Microbiology, 68(8), 2478-2484.

[36] Nouioui, I., Ghodhbane-Gtari, F., Pötter, G., Klenk, H. P., & Goodfellow, M. (2023). Novel species of Frankia, Frankia gtarii sp. nov. and Frankia tisai sp. nov., isolated from a root nodule of Alnus glutinosa. Systematic and Applied Microbiology, 46(1), 126377.

[37] Ramadan, A. M., Nazar, M. A., & Gadallah, N. O. (2021). Metagenomic analysis of rhizosphere bacteria in desert plant Calotropis procera. Geomicrobiology Journal, 38(5), 375-383.

[38] Li, P., Liu, J., Saleem, M., Li, G., Luan, L., Wu, M., & Li, Z. (2022). Reduced chemodiversity suppresses rhizosphere microbiome functioning in the mono-cropped agroecosystems. Microbiome, 10(1), 1-15.

[39] Yue, Y., Shao, T., Long, X., He, T., Gao, X., Zhou, Z., ... & Rengel, Z. (2020). Microbiome structure and function in rhizosphere of Jerusalem artichoke grown in saline land. Science of the Total Environment, 724, 138259.

[40] Zhang, H., Sekiguchi, Y., Hanada, S., Hugenholtz, P., Kim, H., Kamagata, Y., & Nakamura, K. (2003). Gemmatimonas aurantiaca gen. nov., sp. nov., a gram-negative, aerobic, polyphosphate-accumulating micro-organism, the first cultured representative of the new bacterial phylum Gemmatimonadetes phyl. nov. International journal of systematic and evolutionary microbiology, 53(4), 1155-1163.

[41] Wei, C., Gu, W., Tian, R., Xu, F., Han, Y., Ji, Y., ... & Wu, W. (2022). Comparative analysis of the structure and function of rhizosphere microbiome of the Chinese medicinal herb Alisma in different regions. Archives of Microbiology, 204(7), 448.

[42] Hong, S., Jv, H., Lu, M., Wang, B., Zhao, Y., & Ruan, Y. (2020). Significant decline in banana Fusarium wilt disease is associated with soil microbiome reconstruction under chilli pepper-banana rotation. European Journal of Soil Biology, 97, 103154.

[43] Zeng, Y., Nupur, Wu, N., Madsen, A. M., Chen, X., Gardiner, A. T., & Koblížek, M. (2021). Gemmatimonas groenlandica sp. nov. is an aerobic anoxygenic phototroph in the phylum Gemmatimonadetes. Frontiers in microbiology, 11, 606612.

[44] Zeng, Y., Feng, F., Medová, H., Dean, J., & Koblížek, M. (2014). Functional type 2 photosynthetic reaction centers found in the rare bacterial phylum Gemmatimonadetes. Proceedings of the National Academy of Sciences, 111(21), 7795-7800.

[45] Vavourakis, C. D., Andrei, A. S., Mehrshad, M., Ghai, R., Sorokin, D. Y., & Muyzer, G. (2018). A metagenomics roadmap to the uncultured genome diversity in hypersaline soda lake sediments. Microbiome, 6(1), 1-18.

[46] Nemr, R. A., Khalil, M., Sarhan, M. S., Abbas, M., Elsawey, H., Youssef, H. H., ... & Hegazi, N. A. (2020). “In situ similis” culturing of plant microbiota: a novel simulated environmental method based on plant leaf blades as nutritional pads. Frontiers in Microbiology, 11, 454.

[47] Eyre, A. W., Wang, M., Oh, Y., & Dean, R. A. (2019). Identification and characterization of the core rice seed microbiome. Phytobiomes Journal, 3(2), 148-157.

[48] Midha, S., Bansal, K., Sharma, S., Kumar, N., Patil, P. P., Chaudhry, V., & Patil, P. B. (2016). Genomic resource of rice seed associated bacteria. Frontiers in microbiology, 6, 1551.

[49] Jalal, R. S., Sheikh, H. I., Alotaibi, M. T., Shami, A. Y., Ashy, R. A., Baeshen, N. N., ... & Baeshen, M. N. (2022). The Microbiome of Suaeda monoica and Dipterygium glaucum From Southern Corniche (Saudi Arabia) Reveals Different Recruitment Patterns of Bacteria and Archaea. Frontiers in Marine Science, 9, 865834.

[50] Meena, R. S., Vijayakumar, V., Yadav, G. S., & Mitran, T. (2018). Response and interaction of Bradyrhizobium japonicum and arbuscular mycorrhizal fungi in the soybean rhizosphere. Plant Growth Regulation, 84, 207-223.

[51] Floc’h, J. B., Hamel, C., Laterrière, M., Tidemann, B., St-Arnaud, M., & Hijri, M. (2021). Inter-kingdom networks of Canola microbiome reveal Bradyrhizobium as keystone species and underline the importance of bulk soil in microbial studies to enhance Canola production. Microbial ecology, 1-16.

[52] Barelli, L., Waller, A. S., Behie, S. W., & Bidochka, M. J. (2020). Plant microbiome analysis after Metarhizium amendment reveals increases in abundance of plant

growth-promoting organisms and maintenance of disease-suppressive soil. PLoS One, 15(4), e0231150.

[53] Jiang, G., Zhang, Y., Gan, G., Li, W., Wan, W., Jiang, Y., ... & Dini-Andreote, F. (2022). Exploring rhizo-microbiome transplants as a tool for protective plant-microbiome manipulation. ISME Communications, 2(1), 10.

[54] Cernava, T., Chen, X., Krug, L., Li, H., Yang, M., & Berg, G. (2019). The tea leaf microbiome shows specific responses to chemical pesticides and biocontrol applications. Science of the Total Environment, 667, 33-40.

[55] Darby, A. C., Choi, J. H., Wilkes, T., Hughes, M. A., Werren, J. H., Hurst, G. D. D., & Colbourne, J. K. (2010). Characteristics of the genome of Arsenophonus nasoniae, son - killer bacterium of the wasp Nasonia. Insect Molecular Biology, 19, 75-89.

[56] Fay, M., Salazar, J. K., Ramachandran, P., & Stewart, D. (2021). Microbiomes of commercially-available pine nuts and sesame seeds. Plos one, 16(6), e0252605.

[57] de Sousa Lopes, L., Mendes, L. W., Antunes, J. E. L., de Souza Oliveira, L. M., Melo, V. M. M., de Araujo Pereira, A. P., ... & Araujo, A. S. F. (2021). Distinct bacterial community structure and composition along different cowpea producing ecoregions in Northeastern Brazil. Scientific Reports, 11(1), 1-12.

[58] Lee, S. M., Kong, H. G., Song, G. C., & Ryu, C. M. (2021). Disruption of Firmicutes and Actinobacteria abundance in tomato rhizosphere causes the incidence of bacterial wilt disease. The ISME journal, 15(1), 330-347.

[59] Hou, Q., Wang, W., Yang, Y., Hu, J., Bian, C., Jin, L., ... & Xiong, X. (2020). Rhizosphere microbial diversity and community dynamics during potato cultivation. European Journal of Soil Biology, 98, 103176.

[60] Pascual, J., García-López, M., Bills, G. F., & Genilloud, O. (2016). Longimicrobium terrae gen. nov., sp. nov., an oligotrophic bacterium of the under-represented phylum Gemmatimonadetes isolated through a system of miniaturized diffusion chambers. International journal of systematic and evolutionary microbiology, 66(5), 1976-1985.

[61] Zhang, Y., Xu, J., Riera, N., Jin, T., Li, J., & Wang, N. (2017). Huanglongbing impairs the rhizosphere-to-rhizoplane enrichment process of the citrus root-associated microbiome. Microbiome, 5, 1-17.

[62] Jiang, G., Zhang, Y., Gan, G., Li, W., Wan, W., Jiang, Y., ... & Dini-Andreote, F. (2022). Exploring rhizo-microbiome transplants as a tool for protective plant-microbiome manipulation. ISME Communications, 2(1), 10.
